# Saturation mutagenesis and NMR of a fold switch reveal position-dependent deep learning performance

**DOI:** 10.64898/2026.08.02.742283

**Authors:** Nathaniel Felbinger, Kathleen Joyce Carillo, Yihong Chen, John Orban, Brian G. Pierce

## Abstract

A predictive understanding of the properties of fold-switching or metamorphic proteins can provide insights into protein dynamics and energetics, and enable the design of complex protein functions and molecular machines. Recently developed deep learning modeling tools, including AlphaFold, have led to dramatic increases in structural modeling accuracy, but their performance for prediction of fold switching or point mutant effects is unclear. Here we used NMR to measure all substitutions at three key positions of the G_A_/G_B_ fold switch that differentially adopt α-helical (3α), β-grasp (4β+α), or unfolded states, generating a quantitative experimental benchmark for mutation-level fold switching. Using this dataset to assess and compare a panel of deep learning and physics-based modeling and design algorithms, we identified variable and position-dependent performance across methods. Some AlphaFold2-based algorithms were able to predict mutant effects at individual sites, indicating partial sensitivity to physical effects of residue substitutions. Additional comparisons of predictions with experimentally measured stability changes further highlighted position-dependent success and general challenges for predictive algorithms. Together, this study provides a rigorous new benchmark for mutation-induced fold switching and reveals the capabilities and limitations of current deep learning models, establishing a foundational baseline for facilitating the next generation of prediction tools.

## Introduction

In structural biology, proteins are often considered to fold into a single three-dimensional structure determined by their sequence (1). This simplified view fails to capture the true nature of proteins as often dynamic molecules capable of demonstrating a high degree of conformational heterogeneity (2–4). While the magnitude of such conformational transitions and the timescale in which they take place can vary dramatically (5, 6), it is often the case that these dynamics have an important role in the function of proteins. A small-scale vibrational movement in the active site of the isomerase enzyme cyclophilin A can shift and increase its activity within picoseconds (7), whereas large viral glycoproteins such as the SARS-CoV-2 spike protein will shift from a prefusion state to a postfusion state during infection over the course of microseconds or longer (8). Both extremes of this spectrum of conformational flexibility demonstrate that a protein’s ability to shift to alternative states allows it to be more responsive to its environment and participate in dynamic cellular processes.

Fold-switching proteins are capable of undergoing a reversible transition between two distinct stable states, (9, 10), and are a particularly interesting class to study in the context of protein dynamics. A defining property of fold-switching proteins (also referred to as metamorphic proteins) is that movement from one state to another involves a rearrangement of organized secondary structure elements into a distinct and well-ordered fold with a comparable energetic minimum (11). They are distinguished from intrinsically disordered proteins, which can populate multiple states but do not occupy a distinct energetic minimum and lack stable tertiary structure (12, 13). The evolution of naturally occurring fold-switchers that can attain two stable states has implications for protein design. Understanding the fundamental principles will enable the design of proteins that are able to reversibly change between an active and inactive state or multiple distinct functions through predictable modification of environmental conditions.

Accurate prediction of fold-switching properties in proteins, including the structures of the states, remains a major challenge in computational biology (14). Such methods would be useful in expanding upon the limited set of identified natural fold-switching proteins, as well as in the design of novel fold-switching proteins to carry out multiple functions. Previous efforts have utilized sequence-based and structure-based computational methods (15, 16) to aid in the identification of fold-switching proteins from structure in the Protein Data Bank (PDB), leading to a dataset of approximately 100 unique proteins (10). Additionally, recent work has revealed a link between patterns of hydrophobic packing and a temperature-driven switch mechanism (16), suggesting that many metamorphic proteins may be cold-destabilized (17).

Machine learning-based methods to predict protein structure from sequence such as AlphaFold2 (18) and AlphaFold3 (19) have demonstrated high levels of accuracy for various structural classes (20, 21). Despite its impressive performance in structure prediction, AlphaFold has demonstrated limited performance in certain tasks, such as the inability to predict the different states of fold-switching proteins. Previous benchmarking has indicated that AlphaFold captures both states of less than a quarter of documented pre-existing fold-switching proteins (14, 22). To address this, several modifications of AlphaFold have been developed, including AF-Cluster (23), AFSample2 (24), and CF-random (25), which modify the multiple sequence alignment (MSA) input used in predictions to help diversify structural output and improve sampling of alternative states. While able to capture both states of a greater number of fold-switching proteins, the predictive accuracy of some algorithms has been questioned, with some noting a dependence on structures memorized during training (22) rather than a more fundamentally learned energy landscape, while others have disputed that claim (26). An additional limitation of AlphaFold is the poor performance in capturing the structural impact of point mutations (27, 28). While it has been noted that AlphaFold’s high predictive accuracy can be attributed in part to a learned energy function (29), the observed limitations in modeling of alternative states and mutant effects raise questions regarding the extent of its understanding of energetics. Further assessment of AlphaFold and related deep learning approaches in such challenging predictive scenarios would help to address competing interpretations on such trained models and their utility. As noted recently, there has been a lack of available large datasets of NMR-measured mutant conformation (30), which has hindered such analysis.

In this study, we present a systematic assessment of recently developed deep learning modeling methods in capturing the effect of mutations on fold-switching proteins. This evaluation utilizes a new set of point mutations in the G_A_/G_B_ fold switch system (31–34), in which three key positions have been systematically mutated, and each mutant’s folded state(s) characterized by NMR. Previously documented mutants that alter the state equilibrium of the designed metamorphic Sa1 protein (35, 36) and KaiB protein (23) are also included in the assessment. Methods in the evaluation include a range of AlphaFold2-based methods along with other more recently released deep learning methods. While biases were found to be present in evaluated methods, some methods showed limited and position-dependent predictive capabilities for mutant effects, with AlphaFold2 and AlphaFold2-based methods demonstrating surprisingly accurate predictive performance for the population of mutants at residue 45 of the G_A_/G_B_ switch. Overall, our study provides a rigorous new benchmark for mutation-dependent fold switching and defines the capabilities and limitations of current deep learning models, establishing a foundational baseline for developing the next generation of prediction tools.

## Results

### NMR structural characterization of G_A_/G_B_ system mutants

To probe the residue-level dependence of fold switch states, we performed scanning mutagenesis at three key sites in the 56-residue G_A_ (3α) / G_B_ (4β+α) model system (34), measuring fold states using NMR. For each of the three mutation sites in that system, residue L45 in G_A_98, residue T25 in G_B_98, and residue L20 in G_B_98-T25I, we generated all 19 amino acid substitutions and evaluated the relative populations of 3α, 4β+α, and unfolded states using two dimensional ^1^H-^15^N HSQC NMR spectroscopy (**Figure 1, Figures S1-S3**). As G_B_98 only differs from G_A_98 by one residue change (L45Y), the sets of point mutations were performed in nearly the same context. All 60 parent and variant proteins were expressed as soluble monomeric proteins based on their NMR characteristics. No other folded states beyond 3α and 4β+α were detected based on features in the HSQC spectra.

**Figure 1.**
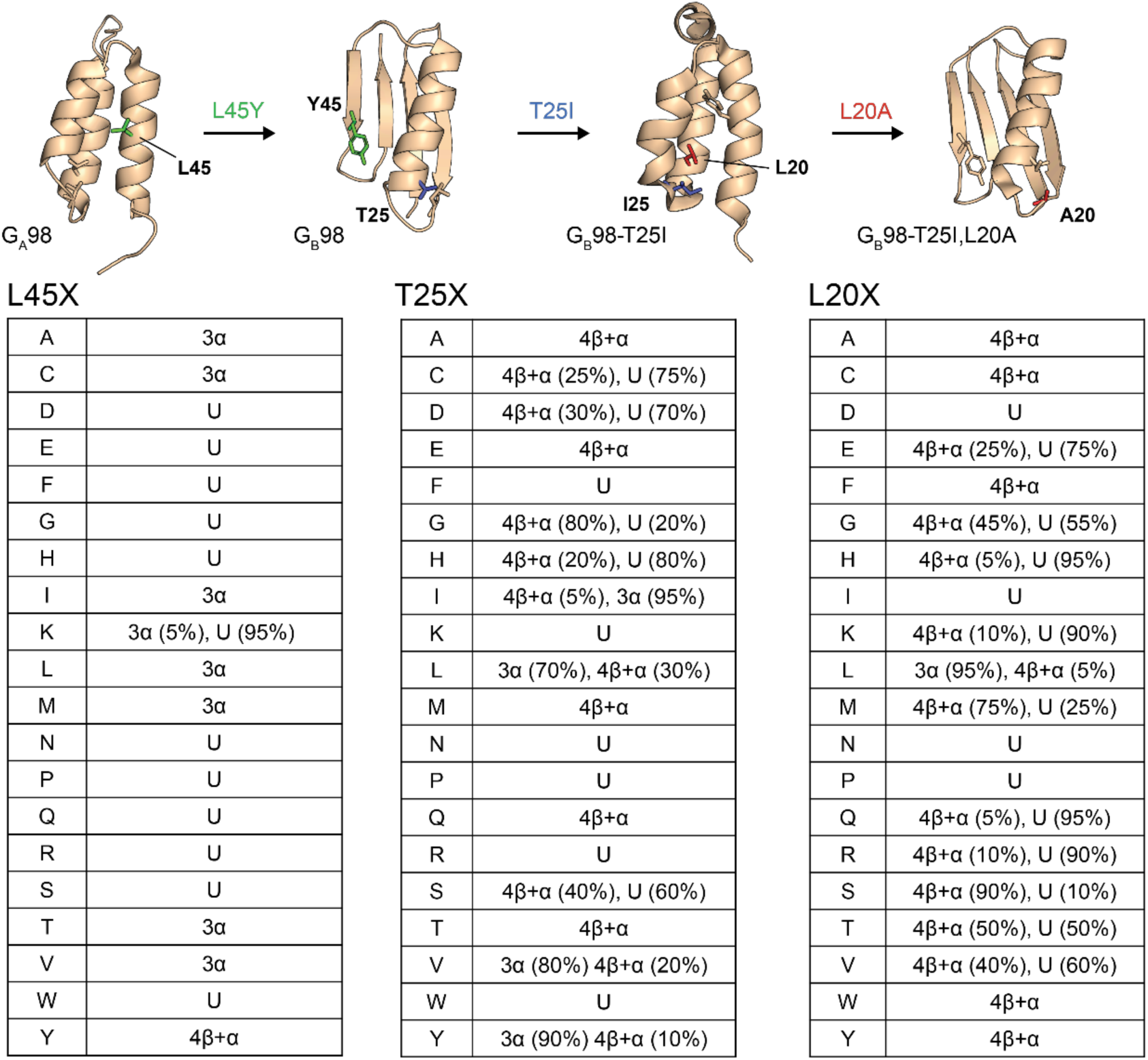
Amino acid dependence of the G_A_/G_B_ 3α/4β+α fold switch. (top) Fold switch states and substitutions leading to transitions, with representative structures (PDB codes 2LHC, 2LHD, 2LHG, and 2LHE) shown from left to right. (bottom) NMR-determined fold switch states for amino acids at residues 45, 25, and 20 are listed by position. As indicated at the top, residue 45 substitutions were tested in the G_A_98 sequence context, residue 25 substitutions were tested in the G_B_98 context (G_A_98+Y45), and residue 20 substitutions were tested in the G_B_98+T25I context. “U” indicates unfolded, and percentages denote relative fold populations in cases of state mixtures.

For the L45X mutations in G_A_98, only L45Y switched the fold from 3α to 4β+α (**Figure S1**). As L45 packs against the hydrophobic core in the 3α fold of G_A_98, mutation to residues with similar apolar character stabilizes the 3α state. Thus, the 3α fold is populated exclusively in 7/20 cases (X=A, C, I, L, M, T, V), underscoring the importance of aliphatic side chains at that position. The remaining 12 mutants (X=D, E, F, G, H, K, N, P, Q, R, S, W) were unfolded, highlighting the destabilizing effects of polar and aromatic residues at that position. One of these, L45K, also populated the 3α state weakly (5%). A requirement for 3α to 4β+α fold switching is that residue substitutions need to both destabilize 3α and stabilize 4β+α. Aromatic residues at position 45, including apolar phenylalanine, are clearly destabilizing to the 3α state, presumably due to their relative rigidity and limited degrees of freedom for packing against the hydrophobic core. At the same time, only tyrosine can stabilize the alternative 4β+α state because it forms a strong hydrogen bond from its side chain hydroxyl to the D47 carboxylate in the β3β4 hairpin (37).

Structural preferences are more evenly distributed for the T25X mutations in G_B_98 (Figure S2). Four mutations at this position (X=I, L, V, Y) populate both the 4β+α and 3α states simultaneously. Of the branched aliphatic residues, switching from 4β+α to 3α is favored in the order I > V > L. The β/γ-branching of ILV at residue 25 likely destabilizes the 4β+α state through steric interactions with the adjacent L20 and D22 side chains, while at the same time stabilizing the alternative 3α state through packing interactions with its hydrophobic core, leading to similar energies for the two states. In this regard, the β-branching of isoleucine and valine appears to be slightly more favorable for 3α formation than the γ-branching of leucine. Interestingly, Y25 has a similar effect to I25, also promoting significant switching to the 3α fold. Notably, T25F and T25W aromatic residue substitutions destabilize 4β+α, most likely through steric clash. Of the aromatic amino acids, only tyrosine can stabilize the alternate 3α state, possibly through side chain hydroxyl-mediated capping interactions with the C-terminus of the α3 helix. In contrast, the 4β+α fold was populated exclusively in 5/20 mutants (X=A, E, M, Q, T) with both apolar and polar residues. Thus, unbranched aliphatics do not promote fold-switching in this sequence context. Another 5/20 mutants (X=C, D, G, H, S) were a mix of 4β+α and unfolded states, and 6/20 mutants (X=F, K, N, P, R, W) were completely unfolded.

For the L20X mutants in G_B_98-T25I, any mutation away from L20 disrupts the 3α state (Figure S3). This is likely because L20 is completely buried in a relatively small 3α hydrophobic core, such that even minor changes from leucine are destabilizing. Five mutants (X=A, C, F, W, Y) switch completely from 3α to 4β+α, while another ten (X=E, G, H, K, M, Q, R, S, T, V) switch to a mix of 4β+α and unfolded states. The remaining four mutants (X=D, I, N, P) are completely unfolded, probably due to a combination of their disruptive effects on the 3α core and the unfavorable steric interactions with the neighboring I25 residue in the 4β+α state.

In summary, the effects on fold switch states vary substantially for each residue position in the respective sequence and structural context. Only one amino acid causes fold-switching in G_A_98-L45X (X=Y), whereas four residues lead to a mixture of 4β+α and 3α states in G_B_98-T25X (X=I, L, V, Y), and fifteen residues induce varying degrees of fold switching and unfolded states in G_B_98-T25I, L20X (X=A, C, E, F, G, H, K, M, Q, R, S, T, V, W, Y).

### Predictive performance of machine learning methods for modeling of G_A_/G_B_ mutants

To assess the capabilities of current deep learning modeling approaches for predicting point mutant effects on fold switch states, we used a set of methods to model all 60 G_A_/G_B_ residue 20, 25, and 45 variants that were characterized by NMR. Assessed methods included AlphaFold-based approaches previously benchmarked in the modeling of fold-switching proteins including AlphaFold2 (18), AlphaFold3 (19), AF-Cluster (23), AFSample2 (24), and CF-Random (25). ESMFold2 (38), ESM3 (39), and Boltz-1x (40) were also included in the analysis to represent deep learning structure prediction methods not based on AlphaFold. All modeling was performed without structural templates, to avoid bias from structures in the PDB.

Sets of models (N=1000 per mutant) produced by each method were analyzed for predicted fold state preference, with each model categorized as 3α (G_A_ fold), 4β+α (G_B_ fold), or neither based on alignment of each model to reference experimental G_A_ and G_B_ structures with TM-align (41). These categorized models were treated as virtual NMR populations where for example, if 700 3α models, 0 4β+α models, and 300 unfolded models are produced for a particular mutant the fold outcome would be defined as 70% 3α, 30% U. To assess each method’s capability to predict effects of mutations at each site, we used the RV coefficient, which is a multivariate version of the Pearson coefficient, comparing the proportion of each fold state of the model outputs to actual fold state preferences determined by NMR (**Figure 2a**). For residue 45 mutants, two methods outperformed the others with highest RV values along with strong significance based on false discovery rate (FDR) permutation testing (q < 0.01): AlphaFold2 (RV = 0.49) and AFSample2 (RV = 0.48). ESM3 had the same significance level with more modest RV (0.22), while CF-Random (RV = 0.37) and AlphaFold2 without MSA (RV = 0.2) showed moderate but still significant FDR levels (q < 0.05). The predictive performance of residue 20 and 25 variants was markedly lower overall than for residue 45, with the majority of tested methods showing lower than 0.2 RV correlation coefficient. Notable exceptions are CF-Random, which had an RV value of 0.38 for residue 25 variants, and AF-Cluster, with an RV value of 0.46 for residue 20 variants; both values are significant based on FDR analysis (q < 0.05).

**Figure 2.**
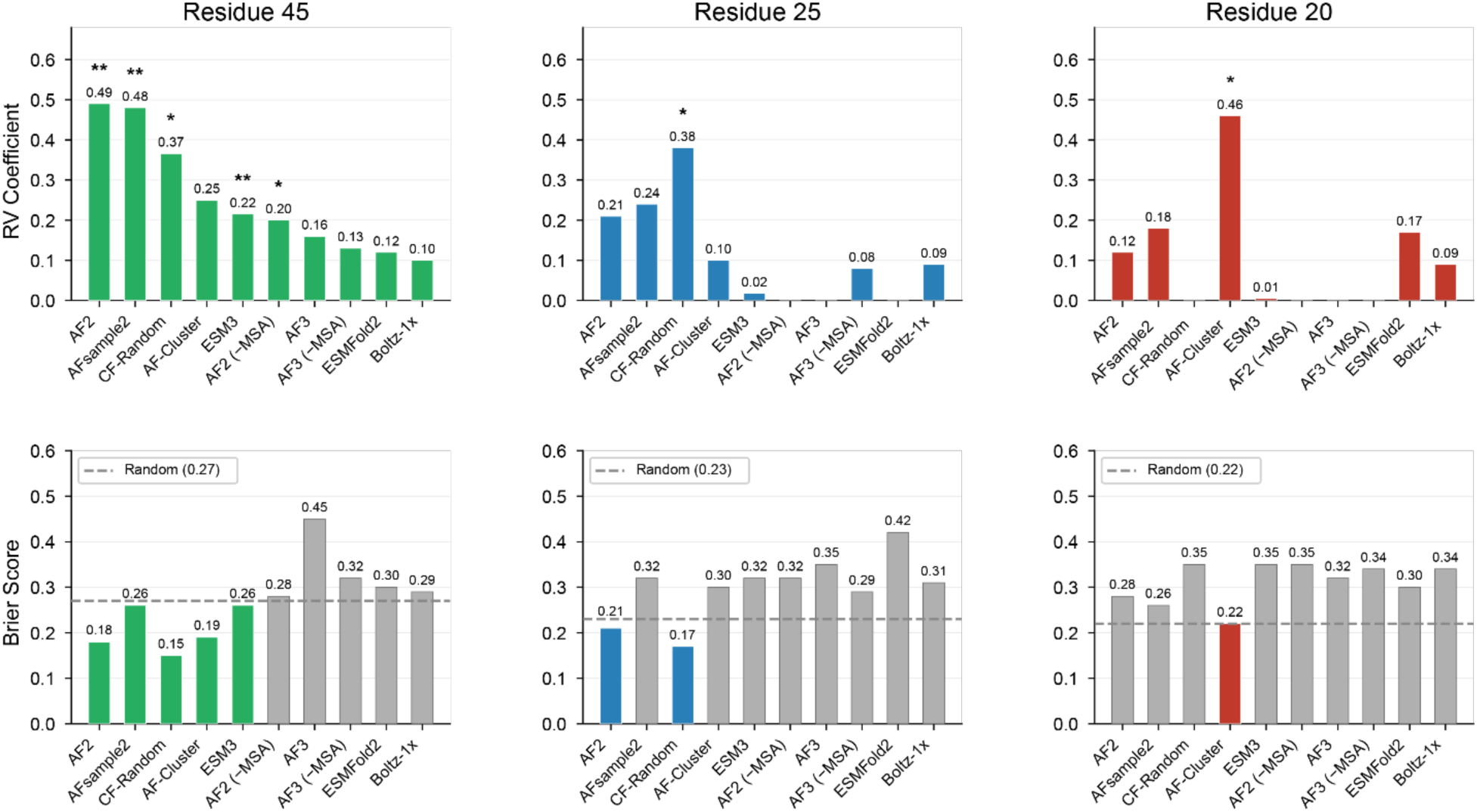
Comparison of deep learning structure prediction methods in capturing effects of mutations on the 3α/4β+α switch. **(a)** All methods were used to produce 1000 models for each mutant at positions 20, 25, and 45 of the G_A_/G_B_ sequences. Models were classified as adopting a 3α fold, 4β+α fold, or being neither based on TM-score alignment to the 3α and 4β+α reference structures (2LHC/2LHD). Proportions of each state calculated from NMR spectra were converted to expected counts out of 1000 structures. For each method, RV coefficients were calculated by the comparison of model proportions to the converted NMR-calculated structure proportions. Absence of a bar indicates the method’s output had no variance. Significance of RV values was assessed with permutation testing, with Benjamini-Hochberg FDR-adjusted significance denoted above methods (* q < 0.05, ** q < 0.01). **(b)** Brier scores were calculated using a similar protocol for quantifying model outcomes and NMR-calculated fold state occupancy. Threshold values for random selection out of the three-class variables were established for each set of mutations individually. Gray bars indicate that the method performed poorer than random baseline.

To quantify the error between predicted fold states and NMR-determined states we calculated Brier scores for predictions at each position (**Figure 2b**). This provides an absolute metric of state prediction accuracy, rather than the prediction of relative states for mutants reflected by the RV coefficient. For residue 45, Brier scores were lowest for CF-Random, AlphaFold2, and AF-Cluster methods, ranging from 0.15-0.19 versus a random baseline of 0.27. AlphaFold2 and CF-Random also showed better than random Brier performance for residue 25 mutants, with scores of 0.21 and 0.17 respectively, versus random baseline of 0.23. For residue 20, although AF-Cluster outperformed the other predictive methods for Brier score (0.22), its score was tied with the random baseline.

Given the indicated residue position-dependent performance of the deep learning methods, we sought to explore the possibility that mutant fold outcomes may also serve as a determinant for predictive performance. Mutants from all positions were pooled together and categorized as either 3α, 4β+α, unfolded, or mixed based on their NMR-determined fold outcomes. Model outcomes were then compared in a pairwise manner. The differences in the number of models produced that belonged to each state were calculated across all tools and all mutants were ranked (**Figure S4**). AUROC values were calculated based on the ability of each ranking to discriminate between the paired experimental fold outcomes. Most tools were found to have stronger performance in discriminating 3α versus Unfolded and 3α versus 4β+α fold outcomes compared to 4β+α versus Unfolded and Mixed versus Uniform mutant populations. This may indicate that predictive capabilities for mutants with a 3α fold outcome is better. However, as the majority of the mutants with a 3α fold outcome are found at position 45 this finding may also still be tied to position dependence.

### AlphaFold2 Predictions of Residue 45 Mutants

To examine the model outcomes of the best-performing predictors of residue 45 mutations, a heat map of model output grouped by NMR-determined fold preference was generated for AlphaFold2 (**Figure 3**). Bias toward the 3α fold is present throughout all 20 mutants regardless of NMR-determined state preference. The average model outputs are distinct between each fold group and appear to correctly correspond to experimental outcomes, in accordance with the relatively high RV and low Brier scores. Model outputs for mutants that fold into the 3α state are on average 96.9% 3α, which is 54.7% and 31.5% higher than the averages of the 4β+α and unfolded mutants, respectively. 45Y, the only exclusive 4β+α folding mutant in the residue 45 set, has a 38% 4β+α model output proportion, 25% higher than any other mutant. To further examine how these fold preferences are reflected across the individual predictions, model TM-scores against the reference 3α and 4β+α were plotted as scatterplots (Figure 4). Mutants 45I, 45Y, and 45P were selected as representative mutants for the 3α, 4β+α, and unfolded groups. All mutants exhibited a dense clustering of high confidence 3α models, once again reinforcing the bias towards the 3α fold. The distribution of models that did not align to either state varied between mutants, with 45Y exhibiting clustering of higher-confidence mutants with secondary structure similar to the 4β+α fold, and 45P producing a wider variety of lower-confidence models with little secondary structure.

**Figure 3.**
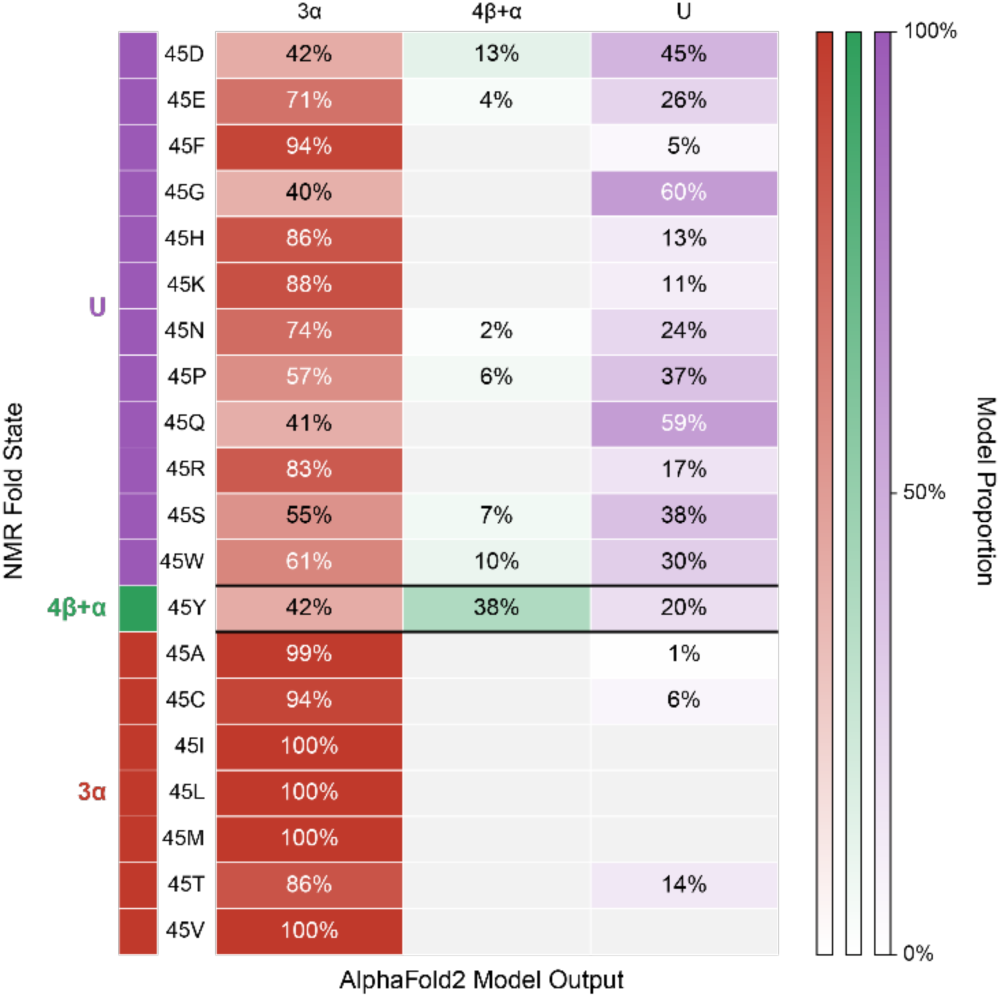
AlphaFold2 demonstrates sensitivity to G_A_/G_B_ residue 45 mutants despite bias for 3α state. AlphaFold2 was used to produce 1000 models of each G_A_/G_B_ mutant at residue position 45. The percentage of models that are in either the 3α fold, 4β+α fold, or neither (denoted as unfolded) are visualized as a heat map. The rows of the graph are comprised of individual mutants grouped by NMR-determined fold propensity. Columns represent the fold outcome proportions of the AlphaFold2 predictions.

**Figure 4.**
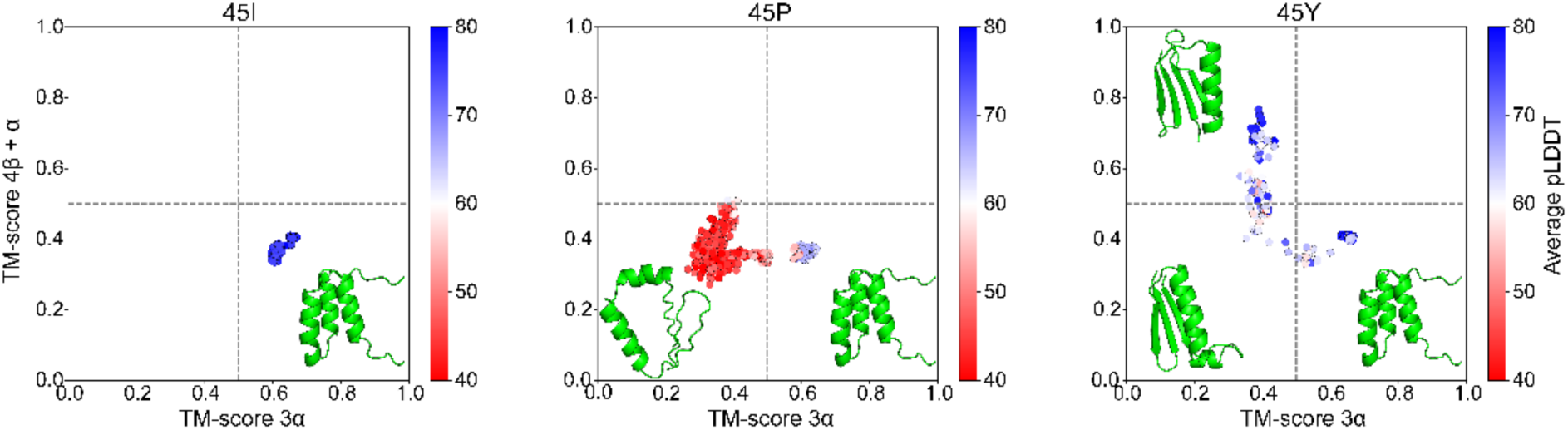
Distribution of output folds from AlphaFold2 predictions of residue 45 mutants corresponds to NMR findings. Individual TM-score outputs of representative 3α fold, 4β+α fold, and unfolded mutants 45I, 45Y, and 45P are displayed as scatter plots. TM-score values are calculated from alignment to 3α reference structures 2LHC (x-axis) and 4β+α reference structure 2LHD (y-axis). Individual points are colored by average pLDDT.

To determine whether the apparent bias towards the 3α state could be altered throughout prediction refinement we investigated the trajectories of the model outputs over AlphaFold2’s recycling steps. AlphaFold2 was modified to output 1000 models after 0, 1, 3, and 20 recycling steps and model outcome proportions were incorporated into a heatmap (**Figure 5**). The proportion of 3α models was highest among models produced with 0 recycling steps regardless of true fold outcome. 92.3% of models across all mutants were classified as 3α without recycling, where it was found to decrease to 67.9% at the default 3 recycling steps and further fall to 60.0% at 20 recycling steps. Without recycling the 3α-folding mutants began with only a marginally higher average proportion of 3α models, 99.3%, compared to unfolding-mutants, with a 3α model proportion of 91.9%. The difference in modeling outcome between the 3α and unfolded mutant groups emerged progressively over the recycling steps. The proportion of 3α models produced by 3α-folding mutants decreased an average of only 7.7% between 0 and 20 recycles, while this proportion fell 46.6% between 0 and 20 recycles for the unfolded mutants. The lone 4β+α-folding mutant 45Y started with a much lower proportion of 3α models, 44.0%, than either of the other groups at 0 recycles.

**Figure 5.**
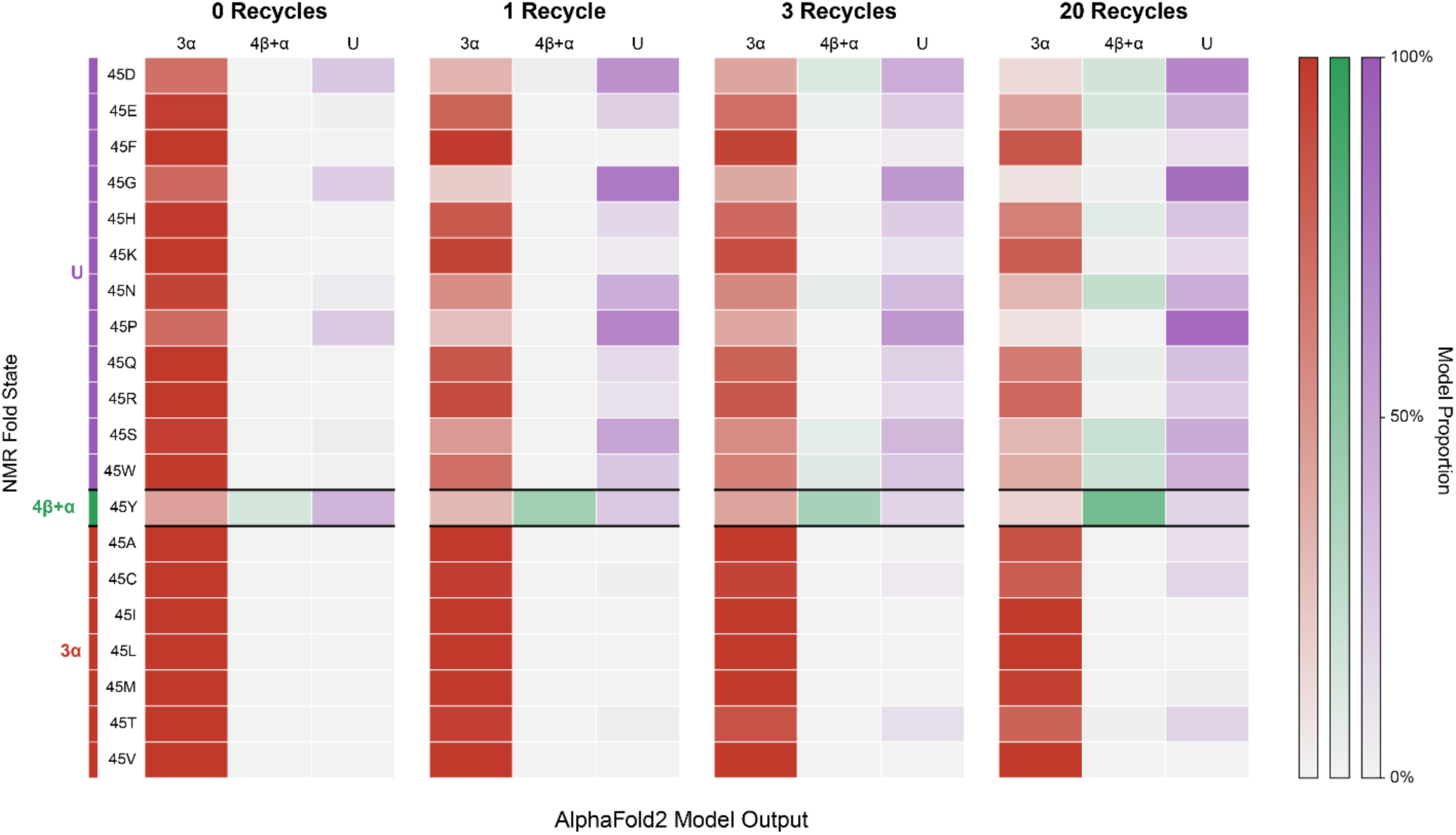
AF2.3 modeling of residue 45 mutants over recycling steps. AlphaFold2 was used to produce 1000 models of each residue 45 variant of the G_A_/G_B_ switch, with configuration adjusted to proceed with 0, 1, 3, and 20 recycling iterations. G_A_/G_B_ fold categorization for each variant is represented, with each row representing a different variant and each column representing the AlphaFold2 modeling outcome proportions.

As the native G_B_-domain sequence contains tyrosine at position 45, it is possible that this propensity to produce more 4β+α models for 45Y may come from association of the 4β+α fold from 45Y-containing structures found in AlphaFold2’s training data. To explore if the presence of tyrosine is responsible for this unique modeling outcome, we modeled two sequences that served as intermediates between the native protein G HSA-binding domain and the designed G_A_ fold switching sequence that contain 45Y. These sequences were previously determined to fold exclusively into the 3α state (42), so the presence of any 4β+α models in the output would indicate memorization from the native G_B_ sequence. However, the model output was composed exclusively of 3α state, indicating that tyrosine at position 45 does not alone drive AlphaFold2 to produce 4β+α predictions (**Table S1**).

With the persistent bias toward the 3α fold, in particular for mutants of residue 45, we also investigated the possibility that homologous proteins with the 3α fold are disproportionately represented in AlphaFold2’s training data. We queried the PDB database for all X-ray and cryo-EM entries with a sequence identity greater than 50% to the G_A_98 sequence released before AlphaFold2.3’s training cutoff date of 09-30-2021. As NMR structures were not used for AlphaFold2 training, they were omitted from the search. 79 hits were found, representing 63 unique protein structures (**Table S2**). These structures were compared to the 3α and 4β+α reference structures using TM-align to assess their fold. It was found that only one of the structures folded into the 3α state, while 46 folded into the 4β+α state and 16 folded into neither state. While the specific training set is not known, these results indicate that a 3α-bias is likely not caused by some form of memorization in the training set.

### Evaluation of physics-based and inverse folding methods

After observing the variable performance of deep learning structural modeling approaches in capturing the impact of our mutants, we assessed the predictive performance of fixed-backbone sequence design approaches, which predict favorable amino acids at each residue position given an input backbone structure. For this effort, we evaluated the predictive capabilities of a panel of inverse folding deep learning methods comprised of ProteinMPNN (43), ESM-IF1 (44), and Frame2Seq (45) as well as a Rosetta (v2.3) physics-based ΔΔG prediction protocol (46). Mutant fold outcomes were classified as 3α, 4β+α, mixed, or unfolded based on NMR measurements. Mutants were considered mixed if they were found to populate multiple states at a level of at least 20% and otherwise classified as their predominant fold state. Representative 3α and 4β+α structures were used as input for mutants of each position: 2LHC/2LHD for residue 45, 2LHG/2LHD for residue 25, and 2LHG/2LHE for residue 20. Each pair comprises the sequence in which mutations were introduced (G_A_98, G_B_98, and G_B_98-T25I, respectively). ΔΔG scores or log probabilities were obtained from respective algorithm output.

Comparison of mutant ΔΔG values predicted by Rosetta with fold outcomes determined by NMR found that Rosetta had some, albeit limited, discriminatory capability for mutant effects with 3α and 4β+α structure inputs (**Figure 6**). Rosetta predicted disruptive effects (> 2 kcal/mol) for most residue 20 mutants with a 4β+α outcome when the 3α structure was used as input, with a neutral predicted effect for these same mutants when the 4β+α structure was input. This suggests that disfavoring the 3α state could be partly responsible for the observed 4β+α state of those mutants. However, residue 20 mutants with mixed-state and unfolded outcomes were predicted by Rosetta to be less disruptive to the 3α state, and unexpectedly, several mutants with unfolded outcomes were not predicted as disruptive in either structural context. For residue 25, as with residue 20, no substantial predicted ΔΔG changes were evident between states when the 4β+α structure was used as the input. When the 3α structure was used as input, mutants with a 3α fold outcome were predicted to be significantly less disruptive than mutants with a mixed fold outcome, but not mutants that cause unfolding. In contrast with the two other fold-switch sites, residue 45 mutants showed disruption to other states when the 4β+α structure was employed as the input, in accordance with the observed fold state changes. However, when the 3α structure was used as input there was no difference in predicted effect of the 3α-folding mutants and mutants that cause unfolding at residue 45. An alternative Rosetta3-based protocol that utilizes an updated energy function and sampling, named Cartesian DDG (47, 48), was also used to predict ΔΔG values for the mutant panel (**Figure S5**). This protocol correctly scored the residue 20 and residue 25 mutants as being less disruptive in the 4β+α state than in the mixed-state and unfolded state. However, it failed to correctly discriminate the fold outcome of any other mutants across the three residue positions. While the incorrect prediction of some effects, including disruptive effects of unfolded mutants, suggests that additional factors need to be considered to fully predict point mutant effects for this challenging multi-state system, physics-based scoring of the 3α and 4β+α static states in Rosetta does seem to agree with a subset of mutant effects at the three switch sites.

**Figure 6.**
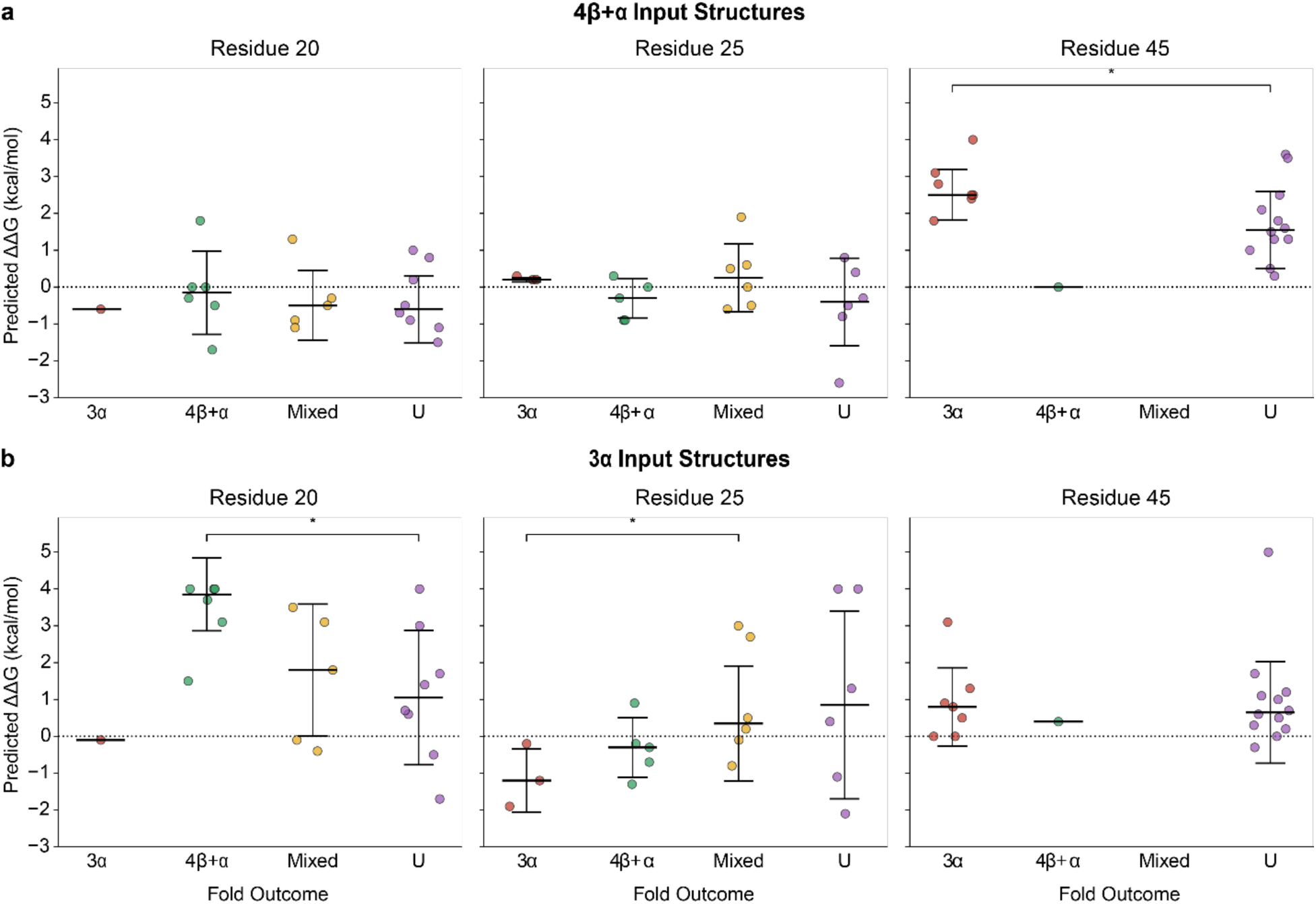
Rosetta ΔΔG predictions of G_A_/G_B_ point mutants. Rosetta v2.3 was used to generate ΔΔG predictions for all mutations at residues 20, 25, and 45 with both **(a)** 4β+α and **(b)** 3α fold reference structures as input. Variants were classified as 3α, 4β+α, or unfolded (U) if they accounted for at least 80% of the estimated fold state based on NMR measurements. If no single fold outcome met this threshold, then the variant was classified as “mixed”. Two-sided Mann-Whitney U tests were used to perform an inter-group comparison between all groups with >1 mutant, and statistically significant comparisons (*, p < 0.05) are indicated above the groups.

Inverse folding log probability scores were likewise assessed for discrimination of different fold states at the three sites with 4β+α (**Figure S6**) and 3α (**Figure S7**) structure inputs. Log probabilities generated from the 4β+α input structures failed to discriminate any of the fold groups from unfolded mutants (**Figure S6**), with minimal overall differences between fold outcomes and tested methods. 3α structure input (**Figure S7**) led to some discrimination between fold outcomes for the inverse folding methods, including at residue 25, as was also seen for Rosetta ΔΔG predictions.

### Mutant fold outcomes in the context of experimental stability

Our G_A_/G_B_ mutants are among the variants measured in the published MegaScale protein folding stability dataset (49), which utilized a high-throughput cDNA display proteolysis method to derive ΔΔG measurements for all point mutants of approximately 500 proteins. To complement our NMR-based structural assessment with a quantitative analysis of folding stability, we assessed computational methods against the mutant ΔΔG values. As ΔΔG only reports the thermodynamics of the unfolding of a single state, there may be limitations to its application in the study of how a protein system interconverts between two stable states. To assess this, we grouped ΔΔG values of mutants reported in the MegaScale set by their NMR preferred fold state to see if unfolding mutants were significantly more destabilizing (**Figure 7**, **Figure S8**). Overall, measured ΔΔG values were clearly higher for NMR-determined unfolded mutants versus mutants with assigned folded or mixed states, corroborating the NMR results. ΔΔG values of unfolded mutants for residue 20 and residue 45 were significantly higher than the folded mutants (p = 0.021 and p < 0.001, respectively), establishing that these measurements were reflective of our structural data (**Figure S8**). No significant difference was observed for the mutants at residue 25, which may be related to the more ambiguous folding outcomes of mutants at this position relative to residues 20 and 45.

**Figure 7.**
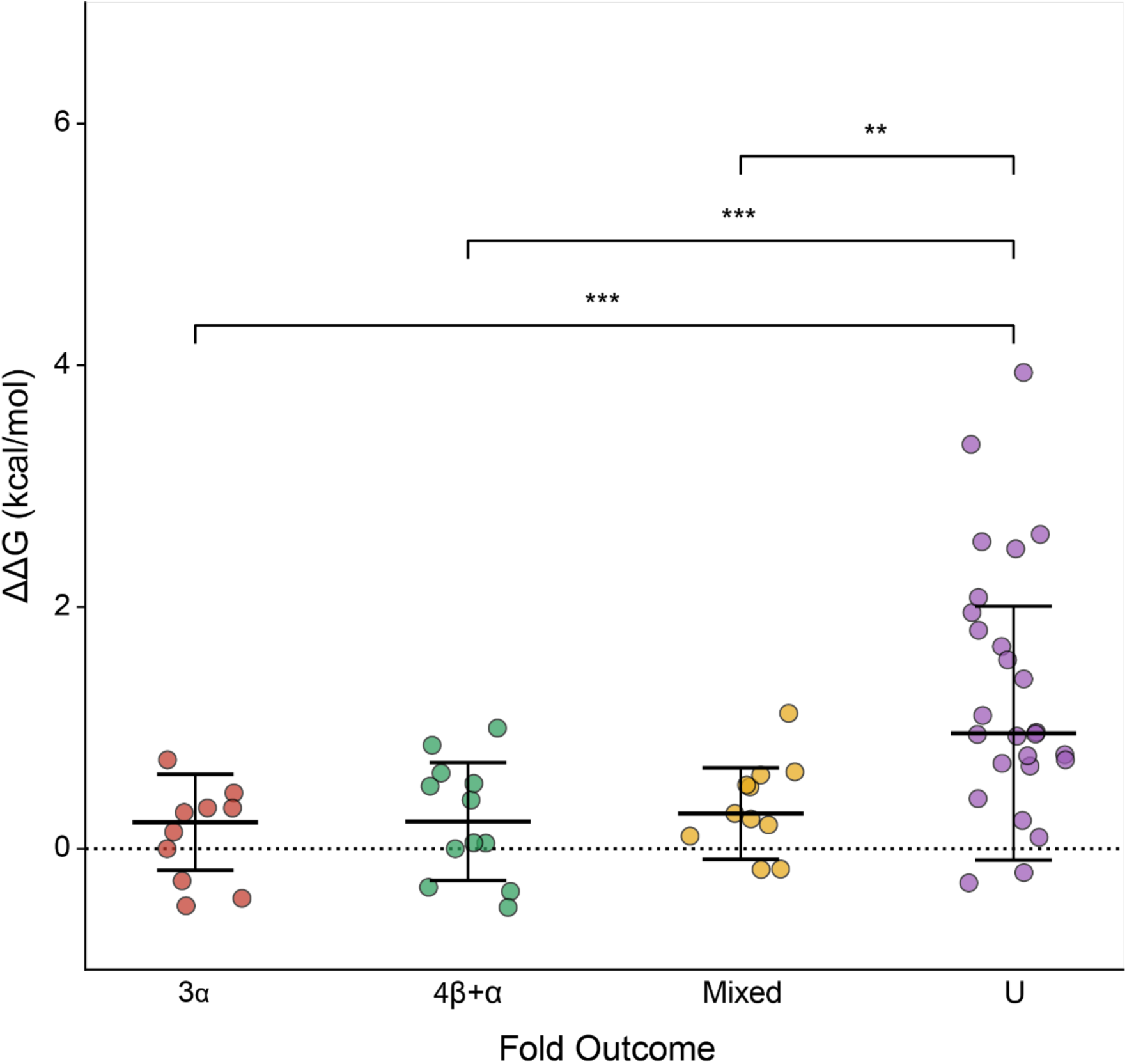
MegaScale stability measurements are associated with NMR-derived fold outcomes. Variants at all three positions were pooled together and grouped by their fold outcomes as measured by NMR spectra. Variants were classified as 3α, 4β+α, or unfolded (U) if they accounted for at least 80% of the estimated fold state. If no single fold outcome met this threshold, then the variant was classified as “mixed”. The MegaScale DMS ΔΔG measurements of each group were compared to themeasurements of the unfolded group using the Mann-Whitney U test (**, p < 0.01; ***, p < 0.001).

The ΔΔG dataset was used to assess whether AlphaFold2 was able to capture the effect that mutations had on folding stability. The proportion of unfolded models produced by AlphaFold2 for each mutant was compared to their respective ΔΔG values. AlphaFold2’s ability to capture the effect of mutations on stability reflected its performance when comparing model outcome to NMR- determined fold preferences. While the proportion of unfolded models had no predictive quality for ΔΔG values of residue 20 mutants (**Figure 8**, r = −0.38, p = 0.109) and residue 25 mutants (r = −0.27, p = 0.250), it was found to strongly correlate with ΔΔG values of residue 45 mutants (r = 0.89, p < 0.001). This would indicate that AlphaFold2 may not only have an understanding of the favored conformation of mutants at this position, but also be able to assess the magnitude of impact the mutations have on the stability of the native folds.

**Figure 8.**
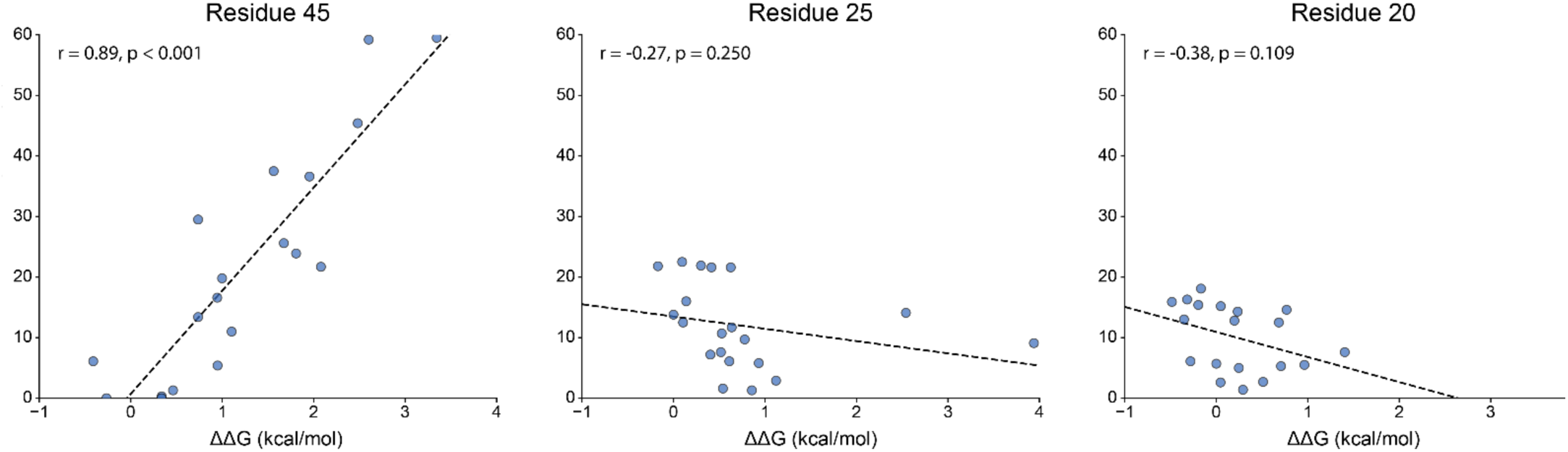
Comparison of AlphaFold2 predictions to MegaScale stability measurements. The proportion of AlphaFold2 predictions classified as unfolded for each variant are compared to their experimentally measured stability change (ΔΔG) values from the MegaScale dataset. Pearson correlation values were calculated for variants of each residue, where a t-test for correlation was used to assess significance of the calculated Pearson correlation values.

This analysis was extended to the previously assessed panel of inverse-folding methods composed of ProteinMPNN, ESM-IF1, and Frame2Seq. Models for both the sequence the mutations are introduced in (2LHC for residue 45, 2LHD for residue 25, 2LHG for residue 20) and models for the outcome state of each mutant (2LHD for residue 45, 2LHG for residue 25, 2LHE for residue 20) were used as input for each method. Log-probabilities for each amino acid were extracted for both sets of structures and separately correlated to experimental ΔΔG values (**Figure S9, Figure S10**). For residue 20, only ESM-IF1 probabilities of the 2LHG G_A_ -fold structure were found to be predictive of ΔΔG values (r= −0.47, p = 0.034). Both Frame2Seq (r = −0.51, p = 0.021) and ESM-IF1 (r = −0.65, p = 0.002) probabilities had moderate predictive character for the effect that residue 25 mutants had on folding stability. Finally, ProteinMPNN and Frame2Seq output probabilities were found to only be predictive of the folding stability impact of mutants of residue 45. The input structures that produce these predictive probabilities differ between the two tools. ProteinMPNN probabilities are predictive of ΔΔG when 2LHC is used as input (r = −0.58, p = 0.012) whereas Frame2Seq demonstrates better predictive capability when 2LHD is used as input (r = −0.46, p = 0.058), though it still does not pass the significance threshold of p < 0.05. When direct comparison of Rosetta predicted ΔΔG values to the MegaScale ΔΔG values were performed, no significant correlations were found with the exception of an incorrect inverse correlation for prediction of residue 25 mutants with the 2LHD input (**Figure S11**).

### Prediction of mutation effects in other fold switch systems

To explore whether these methods demonstrate sensitivity to mutations on other metamorphic systems, 6 NMR-characterized mutants of Sa1 (36) were modeled using AlphaFold2, AlphaFold3, AFSample2, CF-Random, and AF-Cluster. Sa1 is a metamorphic protein designed through the threading of sequences of parent proteins G_A_ and ribosomal protein S6 (35, 36). It is capable of switching between the G_A_ -derived 3α and S6-derived α/β-plait folds where the 3α is favored at lower temperatures but is destabilized and shifted to the α/β-plait fold at higher temperatures. Certain mutations are able to shift the propensity of higher-temperature folding from α/β-plait dominant to a mixed population of α/β-plait and 3α. For our analysis, we selected three mutants that folded primarily (>80%) into the α/β-plait at 25 °C (V88A, V90M, V90T), and three mutants that exhibited a more mixed (∼50/50) population of α/β-plait and 3α states at 25 °C (V90A, V88M, V88T). Mutants were assessed in a similar manner to the G_A_/G_B_ point mutations, where the models produced by each method were compared to a reference structure of each state with TM-align. No method produced any models in the 3α state with the exception of AFSample2, failing to capture the correct 3α-stabilizing effect the mutations had on shifting the state equilibrium (**Figure 9**). The proportion of 3α models output by AFSample2 did not change considerably between mutants and only comprised between 1-3% of all predictions. AF-Cluster did show moderate reductions in the α/β-plait population for the mixed-state mutants, however, with an increase in other/unclassified predictions rather than 3α state.

**Figure 9.**
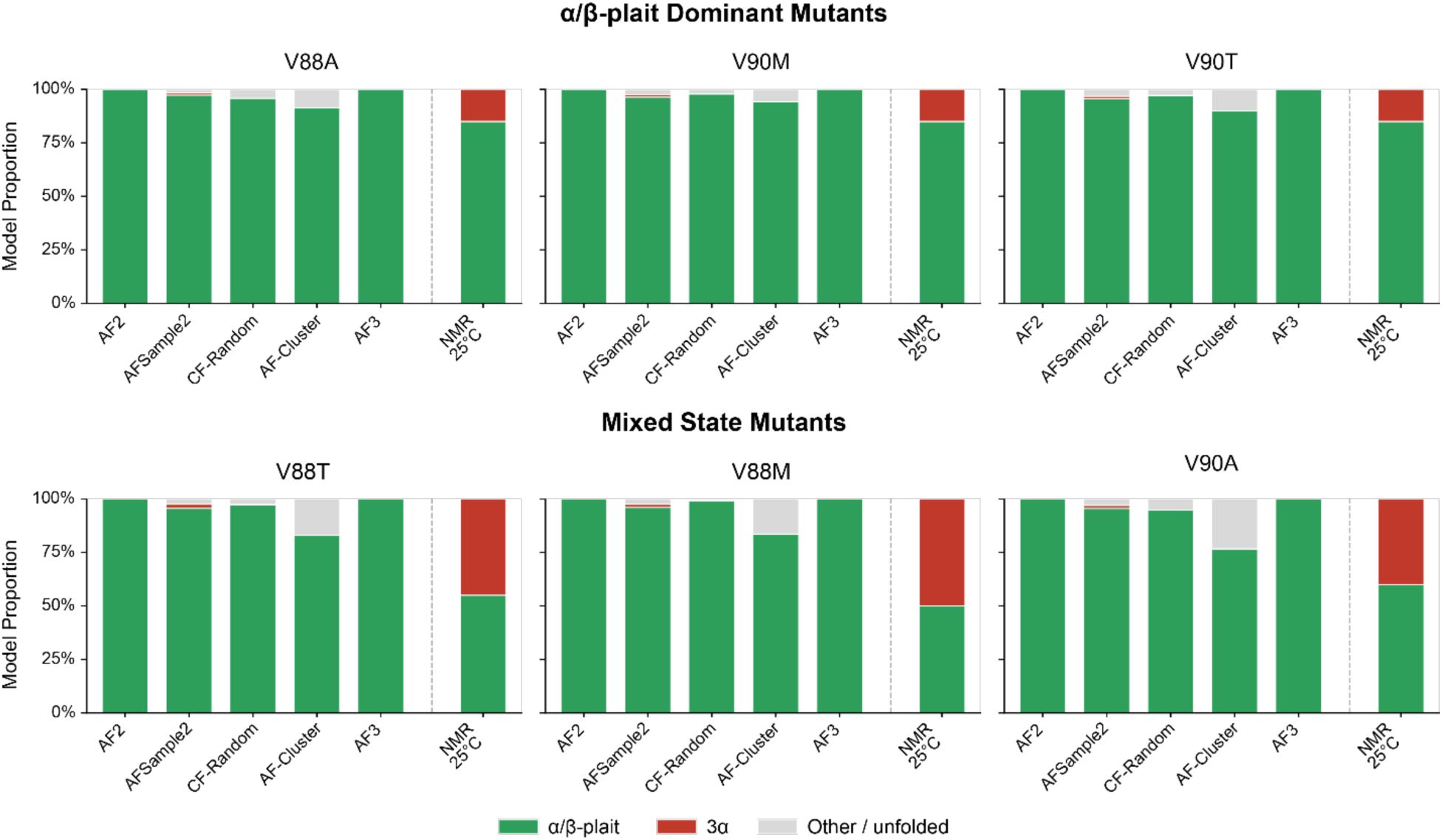
Deep learning structure prediction methods fail to predict the effects of mutations in the Sa1 metamorphic system. AlphaFold2, AlphaFold3, AFSample2, CF-Random, and AF- Cluster were used to produce 1000 models for mutants of the Sa1 metamorphic protein. Model output for Sa1 was classified as being in either the 3α state, α/β-plait state, or neither based on alignment to reference structures 2FS1 and 8E6Y, respectively. A TM-align cutoff of 0.6 was used rather than the 0.5 used for GA/GB, due to higher fold similarity of the two reference state structures. The experimentally determined state proportions at 25 °C are noted in the rightmost column as “NMR 25 °C” for a reference to actual fold outcome.

An additional assessment was performed to study a mutant of the fold switching protein KaiB that was designed to shift the fold equilibrium from a naturally favored ground state to an active fold switched state. The designed sequence consists of three mutations identified with the AF-Cluster method (23) and supported through single-sequence ColabFold modeling. When MSA information was used, all AlphaFold-based methods produced exclusively or near-exclusively fold switch state models for both the WT and designed sequences (**Figure S12**). When MSA information was completely removed, both AlphaFold2 and AlphaFold3 produced exclusively ground state models for the WT sequence and a mix of fold switch state models and unfolded models for the designed sequence, correctly identifying the experimentally determined shift in state favorability, albeit with lower overall proportions of fold switch state for the design than measured by NMR (86%). The AlphaFold2 single sequence (no MSA) result is in accordance with the ColabFold AlphaFold2 modeling of that mutant in the AF-Cluster study (23), while AlphaFold3 was released after that study and thus not tested in that context. The AFSample2 method did not demonstrate any sensitivity to the mutations, while CF-Random did have a small population of ground state models for the WT sequence. Overall, no method that incorporated full or partial MSAs was able to capture the correct fold outcomes of the wildtype and designed mutant sequences.

## Discussion

Our evaluation of computational modeling approaches in the G_A_/G_B_ fold switch system indicates a limited yet measurable ability to recapitulate the effect of mutations on fold state preference. RV coefficient values and Brier scores indicated that predictive performance of modeling methods was generally poor at residue positions 20 and 25 in that system. Conversely, the effect of mutants at residue 45 in G_A_/G_B_ was captured relatively well. We further explored AlphaFold2’s model output for mutants at position 45 and found that, despite an overall bias toward the G_A_ sequence 3α fold, the model output distribution was predictive of experimental outcomes. Physics-based ΔΔG prediction also demonstrated limited performance, appearing to particularly struggle with identifying the destabilizing effect of mutations that cause unfolding from either state. When a subset of the methods was applied to model mutants of the alternative metamorphic protein systems Sa1 and KaiB, there appeared to be little sensitivity to their effect on fold prevalence, particularly for Sa1 variants.

Despite previous work indicating AlphaFold performs poorly in capturing the effect of destabilizing point mutations (27, 28), a recent effort relied on this capability to successfully design a dynamic calcium-binding protein capable of switching between two states (50). Our work similarly supports that AlphaFold does have some limited capacity to capture the effects of point mutations on the conformational outcome of dynamic proteins. It is not entirely clear why such sensitivity to point mutations in the G_A_/G_B_ system was primarily limited to residue 45. In the context of the designed G_A_/G_B_ system, residues 20 and 45 are both components of the hydrophobic core of the G_A_ helical bundle and G_B_ β-grasp folds (42, 51, 52). Neither has been noted as having a more significant impact on stability than the other. However, 45Y is highly conserved among natural G_B_ domains and has been noted as a key contributor to stabilizing the 4β+α fold of the wild-type sequence (42). Although we found that AlphaFold produced exclusively 3α models for previously characterized G_A_ -folding mutants that contain 45Y, MSA-based methods might get particularly strong signal from the conserved residue 45 position.

It is possible that the elevated performance of the residue 45 mutants may not be tied to the position itself. Mutations at the other two residue positions are point-mutants of the G_B_98 sequence rather than the lone drivers of the switch. This means that in the context of GA, the T25X and L20X mutants are effectively more complex sets of double and triple mutants. Another possible explanation is that improved performance is tied to better predictive capability of the 3α state itself. The effect of individual residues on the helix propensity is relatively strict compared to residues that are proximal to coil or turn regions (53, 54). Because of this, the helical geometry of the 3α fold may make it easier to determine the effect of mutations. However, the majority of the 3α-folding mutants are found at residue 45 and it has the lowest prevalence of mixed-state mutants, which is a state class that challenged predictive methods. Our findings related to state-preference of mutants cannot therefore be completely decoupled from possible position-dependence.

While modification of MSA information may help to uncover alternative states for dynamic proteins (23, 24), our results indicate that this does not help increase AlphaFold’s sensitivity to the effect of mutations. These findings support previous studies which indicated that coevolutionary signal from shallow or clustered MSAs may contribute little to the model outcome of AlphaFold2 predictions (22, 55). A previous study assessed the ability of an AlphaFold2-based tool called Cfold to capture the fold outcomes of a limited set of three G_A_/G_B_ mutants, where it was found Cfold produced models of both states regardless of the true fold outcome (56). While AF-Cluster demonstrated some ability to discriminate true fold outcome of a set of 12 G_A_/G_B_ mutants with one or more substitutions based on model confidence, the method still incorrectly produced models in both states for most mutants (23). It is important to note that G_A_/G_B_ is a designed system that is manually tuned to attain the capacity to switch folds. Although it has a relatively high degree of sequence similarity to its natural parent sequence, this may still limit the utility of altering evolutionary signal from an MSA constructed from natural proteins, in accordance with recent work which states that AF-Cluster is not expected to perform well with engineered versus natural proteins (26). Methods such as AF-Cluster have demonstrated the ability to design mutants of natural fold switch proteins such as KaiB that shift fold preference to a particular target state (23). Alternative approaches may need to be developed to better take advantage of and potentially improve this limited capacity to capture the effect of mutations on fold outcome. Tools such as AF2Rave (57) and AlphaFlow (58) have demonstrated success in capturing physically feasible conformations of flexible protein regions by integrating AlphaFold2 predictions into more extensive predictive pipelines. AF2Rave utilizes reduced-MSA AlphaFold2 predictions similar to methods evaluated in this benchmark. However, these predictions serve only as an initial ensemble to initialize an extensive predictive pipeline that integrates enhanced sampling and molecular dynamics approaches. AlphaFlow tasks AlphaFold2 with sampling from a diverse distribution of feasible 3D conformations rather than producing predictions from scratch. However, compared to the secondary structure remodeling of a metamorphic protein, such alternative conformations are relatively subtle, and these approaches have not been assessed in the context of point mutations or fold switch proteins. An additional approach is the fine-tuning or training of open-source models (59) on NMR conformers. Most current machine learning structure-prediction methods, with the exception of Boltz-2 (60), do not incorporate NMR structures in their training sets. With NMR structures being overly represented in the established fold switching dataset compared to all structures in the PDB (10), their exclusion during training may cause suboptimal performance in the modeling of metamorphic proteins and their mutants.

It is possible that AlphaFold’s sensitivity to mutations may not be sufficient to capture the effects of point mutations in the fold of metamorphic proteins and that alternative approaches should be explored for related design efforts (61). Traditional molecular dynamics simulations may be of use to aid in the design of fold switching proteins. However, their computational expense and potential to get trapped in local energy minima (62, 63) may limit their application in designing fold switching protein systems that can undergo extensive secondary structure remodeling over seconds to hours (64). It may be worth exploring the use of alternative MD simulation approaches such as enhanced-sampling methods or coarse-grained simulations which are capable of simulating longer time-scales at a much lower computational cost (65). Methods capable of being fine-tuned on thermodynamic data such as BioEmu (66) may also prove useful in the application of fold switch system design. Continued development of methods that directly incorporate conformational ensembles and thermodynamic data could be productive in improving the design of fold switching and other dynamic protein systems.

Inverse folding methods may also warrant further exploration in this context. Others have recently used inverse folding models to analyze and design variants with altered conformations (30) and to design a de novo Ca^2+^-activated two-state system (50). These methods carry relatively low computational cost compared to deep learning structure prediction methods and MD simulations. Further, these design methods are conditioned on structure rather than sequence. Scoring two structure inputs with an inverse folding method is a more constrained task than having a structure prediction tool predict alternative folds of a given sequence. Our evaluation of representative inverse folding methods revealed poor capability in predicting the fold outcome of mutants. However, we also observed significant correlations between inverse folding scores and MegaScale ΔΔG values for mutants at all three residue positions, indicating some grasp of the energetics of G_A_/G_B._ Previous studies have used other formulations in their inverse folding-based designs of fold switch proteins, such as contrastive scoring (30) or joint optimization (67) to compare scores for a given mutant across different folds. Adapting such formulations, or perhaps fine-tuning inverse folding models specifically on fold switching proteins, may improve their capacity to capture the effects of mutation on fold switching.

This work provides a novel dataset of point mutants in the G_A_/G_B_ fold switch system with mutation-induced effects on the conformational state that are experimentally defined by NMR. Benchmarking of computational tools highlights ongoing challenges and variability between predictive approaches, with some current machine learning structure-prediction methods demonstrating limited sensitivity to these unseen mutants that is likely not from structural memorization. However, MSA-modified protocols that perform relatively well for certain positions or systems do not appear to consistently perform well in other scenarios, highlighting a lack of general utility. Future methods that incorporate information from structural ensembles or more directly incorporate thermostability data during training may better capture mutation-induced changes in the conformational preference of dynamic systems such as G_A_/G_B_.

## Methods

### Protein expression and purification

G_A_98 and G_B_98 mutants were prepared using the QuikChange (Agilent) mutagenesis kit. All mutants were then cloned into a vector that features an N-terminal subtilisin-prodomain fusion tag system (Profinity eXact, Bio-Rad). BL21(DE3) strain *E. coli* cells were transformed with the vector and cultured in M9 minimal media for ^15^N labeling. The cells grew at 37 °C until they reached an optical density (OD_600_) of 0.6-0.8. To induce protein expression, 1 mM IPTG was added, and the culture was incubated overnight at 25 °C.

After overnight incubation, cells were harvested by centrifugation, resuspended in 100 mM potassium phosphate (KPi) buffer (pH 7.0), and lysed by sonication. The cleared cell lysate was injected onto a 5-mL eXact column at a flow rate of 5 mL/min. After sample injection, the column was extensively washed with 100 mM KPi buffer (pH 7.0). The target protein was cleaved and eluted by injecting 6 mL of 10 mM sodium azide in 100 mM KPi (pH 7.0) at a flow rate of 0.5 mL/min. The purity of the protein samples was determined using SDS-polyacrylamide gel electrophoresis. The purified protein samples were then pooled and concentrated for NMR analysis.

### NMR spectroscopy

^15^N isotope-labeled protein samples were prepared and concentrated to a final concentration of 0.05 to 0.3 mM for NMR analysis in 100 mM potassium phosphate buffer (pH 7.0) containing 5% D_2_O. All 2D ^1^H-^15^N HSQC NMR experiments were done at 5°C on a Bruker AVANCE 600 MHz spectrometer with a cryoprobe. Spectra were processed and analyzed using Topspin 4.5.0.

### NMR data analysis

Peaks corresponding to 3α (^1^H 8.5-10 ppm and ^15^N 121.5-131 ppm), 4β+α (^1^H 8.4-9.1 ppm and ^15^N 123-128 ppm), and unfolded state (^1^H 7.5-8.4 ppm and ^15^N 106-130 ppm) were identified. Peaks selected for each state were ensured to be isolated and not overlapping. The peak volumes of the residues corresponding to the 3α, 4β+α, and unfolded state (*V*_α,ave_, *V*_4β+α,ave_, *V*_U,ave_) were determined and integrated. The fractional population of each state was then calculated by dividing *V*_α, ave_, *V*_4β+α, ave_, and *V*_U, ave_ by the total peak volume (*V*_total_), where the *V*_total_ is the sum of the peak volumes for all three states. The calculated fractional populations were then expressed as percentages by multiplying them by 100. The estimated error in the peak intensity measurements is ±5% based on background noise.

### Model generation

#### AlphaFold2

All models were predicted using AlphaFold v2.3.2 (18) run locally, downloaded from its GitHub repository: https://github.com/google-deepmind/alphafold. Predictions were produced with and without the use of MSAs, with both prediction pipelines avoiding the use of structure templates. When used, MSA features were generated individually for each of the 60 predicted mutants with default genetic databases as described in the AlphaFold GitHub setup. When predicting without MSAs, models were generated with the use_precomputed_msas flag active where defined MSA inputs contained only the mutant sequence itself. The five default AlphaFold monomer models were used to produce 200 models each for a total of 1000 output models.

#### AFSample2

AFSample2 (24) version 1.0 was downloaded from Github (https://github.com/wallnerlab/AFsample2). MSAs produced during the vanilla AlphaFold2 runs were reused for AFSample2 predictions. Dropout was enabled for all predictions with vertical MSA randomization being set to 0.1 (10%). All five monomer and monomer_ptm models were used to produce 100 models each, for a total of 1000 output models.

#### AF-Cluster

Models were produced according to the protocol described to evaluate AF-Cluster (23) on previously published G_A_/G_B_ mutants using the Colab page https://github.com/HWaymentSteele/AF_Cluster/blob/main/data_sep2022/colabfold_outputs/GAGB_mutations_in_colabdesign.ipynb. Different mutant sequences produced various numbers of clusters, so the number of seeds used for each mutant sequence was dynamically increased until at least 1000 models were produced.

#### CF-random

CF-random (25) was retrieved from https://github.com/ncbi/CF-random_software. Predictions were generated using the default blind mode as described in the associated GitHub page. 1000 models were produced for each mutant sequence. *AlphaFold3.* Predictions were generated using AlphaFold v3.0.1 (19) available at https://github.com/google-deepmind/alphafold3 with model parameters received from Google. Like AlphaFold2, prediction pipelines both using and not using MSAs were run with both pipelines excluding the use of structural templates. 1000 total models were produced for each mutant both with and without MSAs using seed numbers 1-200.

#### ESM3

Models were generated using v3.1.5 of ESM3 (39) available at https://github.com/evolutionaryscale/esm/releases with ESM3-open model weights downloaded from https://huggingface.co/EvolutionaryScale/esm3-sm-open-v1. Configuration settings were left at their default values for the structural generation track.

#### ESMFold2

Models were generated using a local installation of ESMFold2 (38), downloaded from its GitHub repository: https://github.com/Biohub/esm. The ESMFold2-fast model was used, which does not utilize MSAs in its prediction. 1000 models were produced using seeds 1-1000 with default settings utilized for all predictions.

#### Boltz-1x

Boltz-1x (40) was run using version v1.0.0 which is available at https://github.com/jwohlwend/boltz/releases. The –use_msa_server flag was used to build individual MSA inputs for each mutant, with remaining prediction parameters being left as their default values (step_scale = 1.678, recycling_steps = 3, sampling_steps = 200). 1000 total models were produced for each mutant using 1000 individual sampling runs.

### Inverse folding methods

#### ProteinMPNN

ProteinMPNN (43) was retrieved from the repository at https://github.com/dauparas/ProteinMPNN. The first NMR conformer of the four reference PDBs (2LHC, 2LHD, 2LHE, 2LHG) was used as input with the default v_48_020 full protein backbone model. To retrieve log probabilities, the –unconditional_probs_only flag was active during execution.

#### ESM-IF1

ESM-IF1 v1.0.2 (44) was downloaded from https://github.com/facebookresearch/esm. The first NMR conformer of the four reference PDBs was used as input with the –nogpu and -write_logits flags active.

#### Frame2Seq

Frame2Seq (45) was downloaded from https://github.com/dakpinaroglu/Frame2seq. By default, Frame2Seq will only print the negative-log-likelihood value for the designed residue at each position with the save_indiv_neg_pll flag. To retrieve log probabilities for each residue at each position modifications were made to the Frame2seqRunner.py and design.py files. Specifically, the pred_seq_unscaled tensor containing the full set of residue probabilities was copied, moved to CPU, and log-transformed using the PyTorch package torch.log function.

### Rosetta ΔΔG prediction

ΔΔG predictions were produced using the Rosetta2.3 ΔΔG interface scanning protocol (46) with the monomer flag passed in. Backbones were fixed, with chi angle refinement (-chi) and side-chain repacking with extra rotamers (-ex1 -ex2 -ex3 -extrachi_cutoff 1) carried out for each mutant. As with the inverse folding methods, the first conformer of each representative NMR structure (2LHC, 2LHD, 2LHE, 2LHG) was used as input. Rosetta3 (v3.14) ΔΔG predictions were generated with the cartddg2020protocol (47, 48), using the RosettaDDGPrediction wrapper (48) downloaded from its Github repository (https://github.com/ELELAB/RosettaDDGPrediction). Predictions were made using the default cartddg2020_ref2015.yaml protocol included in the repository.

### Model classification and evaluation

TM-scores calculated using TM-align against pairs of reference structures were used to classify the fold state of model outputs. A TM-score cutoff of 0.5 was used to classify G_A_/G_B_ mutants, with models failing to meet this threshold for either reference structure defined as “neither”. When evaluating the Sa1 and KaiB fold switch systems it was found that some models met this score cutoff for both reference structures, likely due to the smaller scale of structural reorganization between the two reference states. Therefore, for those systems the classification threshold was raised to a TM-score of 0.6. The following PDB entries served as reference structures: 2LHC and 2LHD for GA/GB, 2FS1 and 8E6Y for Sa1, and 2QKE and 5JYT for KaiB. For NMR-determined PDB structures with multiple conformers, the first conformer was used for assessments.

RV coefficients were calculated using the hyppo Python package comparing the model fold output of each mutant to observed NMR structure populations. As 1000 models were produced for each mutant, proportions of each state observed from experimental NMR data were extrapolated to 1000 structures. For example, if 50% 3α state, 0% 4β+α state, and 50% unfolded were observed with NMR, the actual model outcome would be defined as 500 models in the 3α state, 0 in the 4β+α state, and 500 unfolded. Calculation of statistical significance of RV coefficients followed the procedure described in the hyppo.tools.perm_test. Fold outcomes for each method and mutant position were randomly permuted 100,000 times to generate a null distribution. Resulting p-values were corrected for false-discovery rate using the Benjamini-Hochberg procedure, yielding adjusted q-values. Brier scores were calculated using the Scikit-learn Python package’s brier_score_loss function. The fraction of models produced in the 3α state, 4β+α state, or unfolded (neither state) by each method were used as predicted probabilities. These were compared to the fraction of each state observed with NMR. Scores were averaged across all mutants at each residue for each method, with lower scores indicating better agreement between the model fractions produced by the methods and the NMR-determined state population fractions. An equal-probability prediction across the three state categories for each residue position was used as a random baseline.

### Plot and figure generation

Figures of structures and models were generated using PyMOL version 2.5 (Schrodinger, LLC). All plots were generated using the pandas and matplotlib python packages.

## Acknowledgements

This work was supported by National Institutes of Health grants R35GM144083 (to B.G.P.) and R01GM062154 (to J.O.), and grant 1001530 from the W.M. Keck Foundation (to J.O.). Computing resources from the University of Maryland Zaratan and Institute for Bioscience and Biotechnology Research high performance computing clusters were used for this study. The NMR facility is supported by the University of Maryland, the National Institute of Standards and Technology, and the W. M. Keck Foundation.

## Data Availability

All data are available in the figures, supplementary figures, or downloadable from https://doi.org/10.5281/zenodo.22310587. Scripts used for analysis, modified inverse folding code to capture output values, and individual model data are available from the authors upon request.

