## Supplemental Figures and Tables for "Saturation mutagenesis and NMR of a fold switch reveal position-dependent deep learning performance"

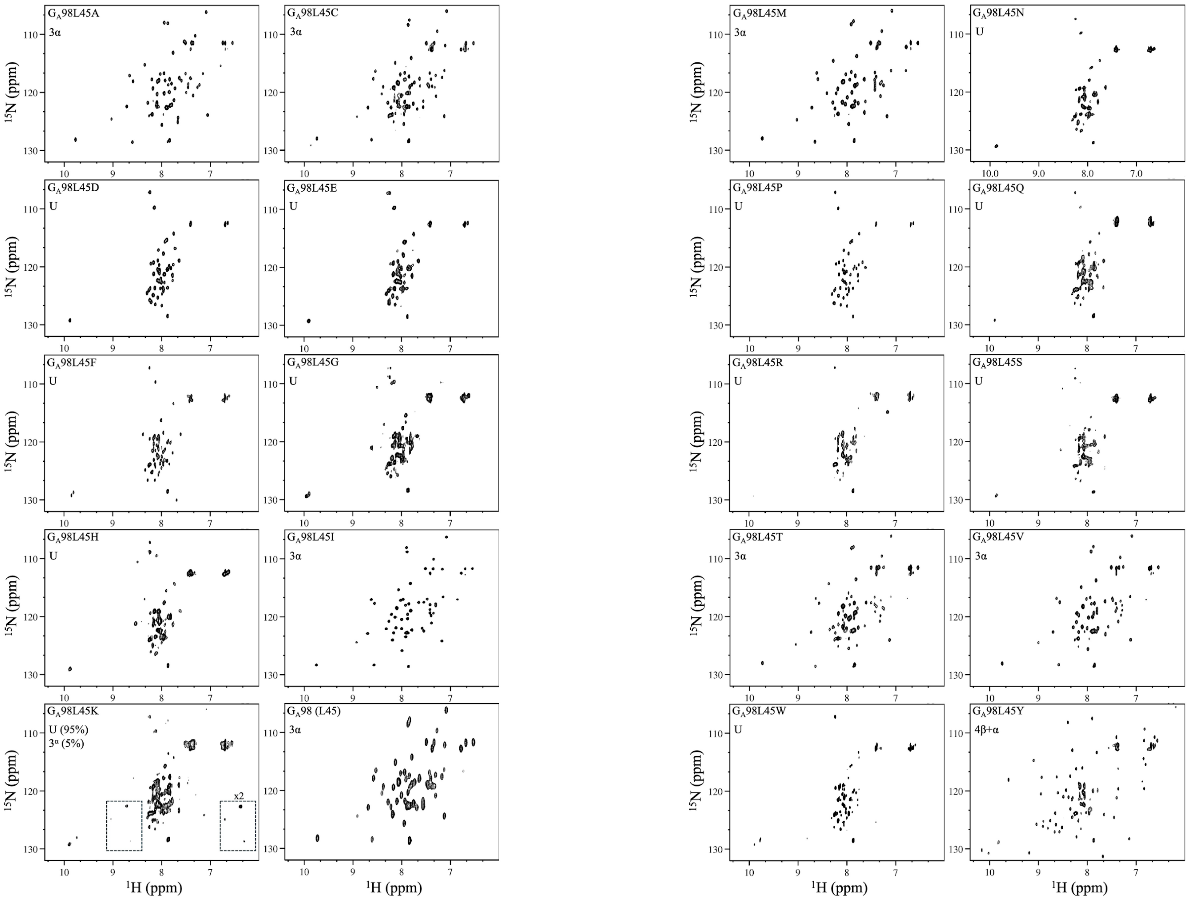


**Figure S1. Two dimensional $^{1}$H-$^{15}$N HSQC spectra for each GA98-L45X mutant.** Low intensity diagnostic signals are highlighted in dashed boxes.

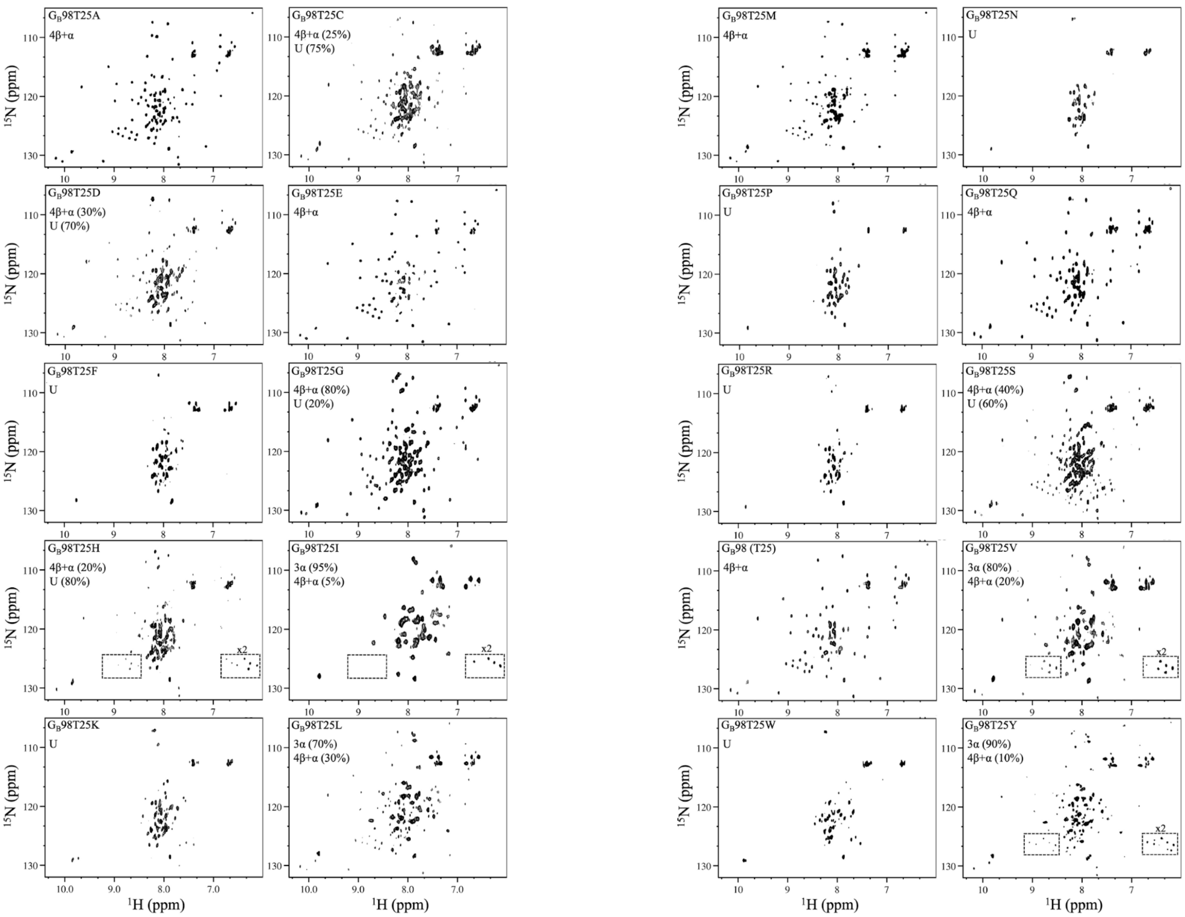


**Figure S2. Two dimensional $^{1}$H-$^{15}$N HSQC spectra for each GB98-T25X mutant.** Low intensity diagnostic signals are highlighted in dashed boxes.

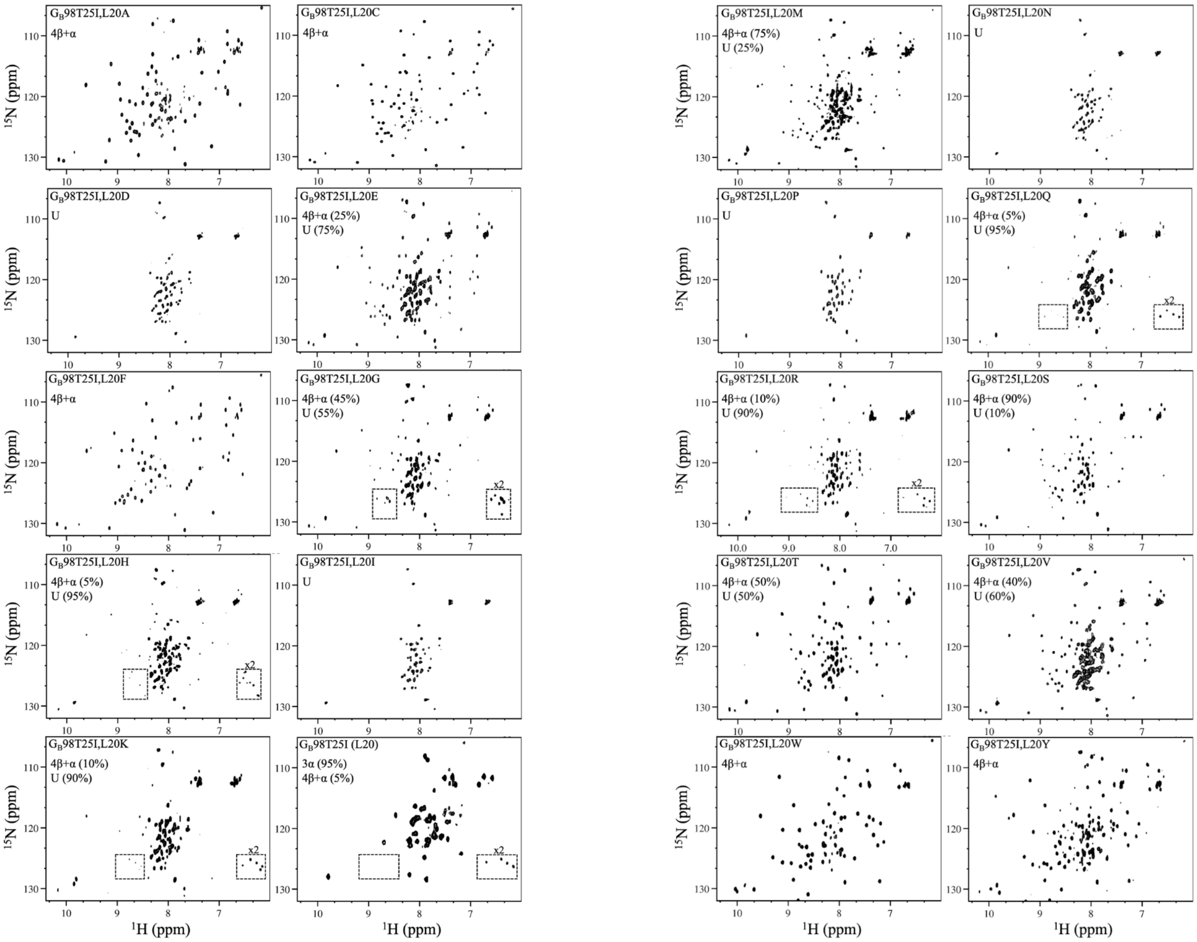


**Figure S3. Two dimensional $^{1}$H-$^{15}$N HSQC spectra for each GB98-T25I, L20X mutant.** Low intensity diagnostic signals are highlighted in dashed boxes.

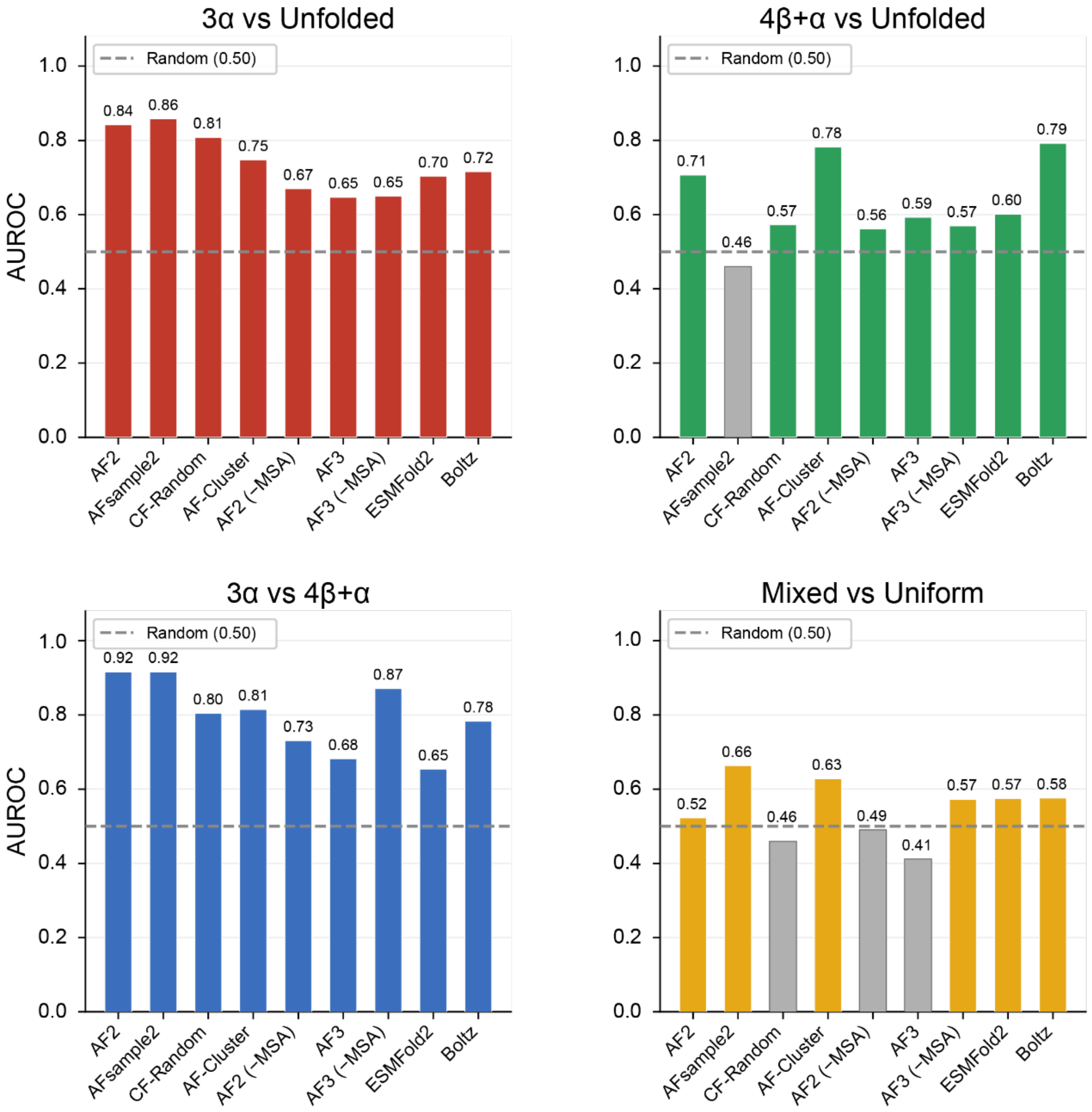


**Figure S4. Pairwise classification performance of fold state outcomes.** Mutants from all positions were pooled together and classified as either 3α, 4β+α, mixed or unfolded based on NMR analysis. If a mutant had an 80% or higher preference for a fold it was classified as being in that state, otherwise it was classified as mixed. For each method, the difference in number of models produced that were in either of the states being compared was calculated. For the mixed vs uniform state comparison, this was instead calculated as 1 – Max (Proportion 3α, Proportion 4β+α, Proportion Unfolded). The mutants were ranked based on these difference values and AUROC calculations were performed. Line for random AUROC baseline (0.5) is displayed as a grey dashed line. Bars of methods below random baseline are colored grey.

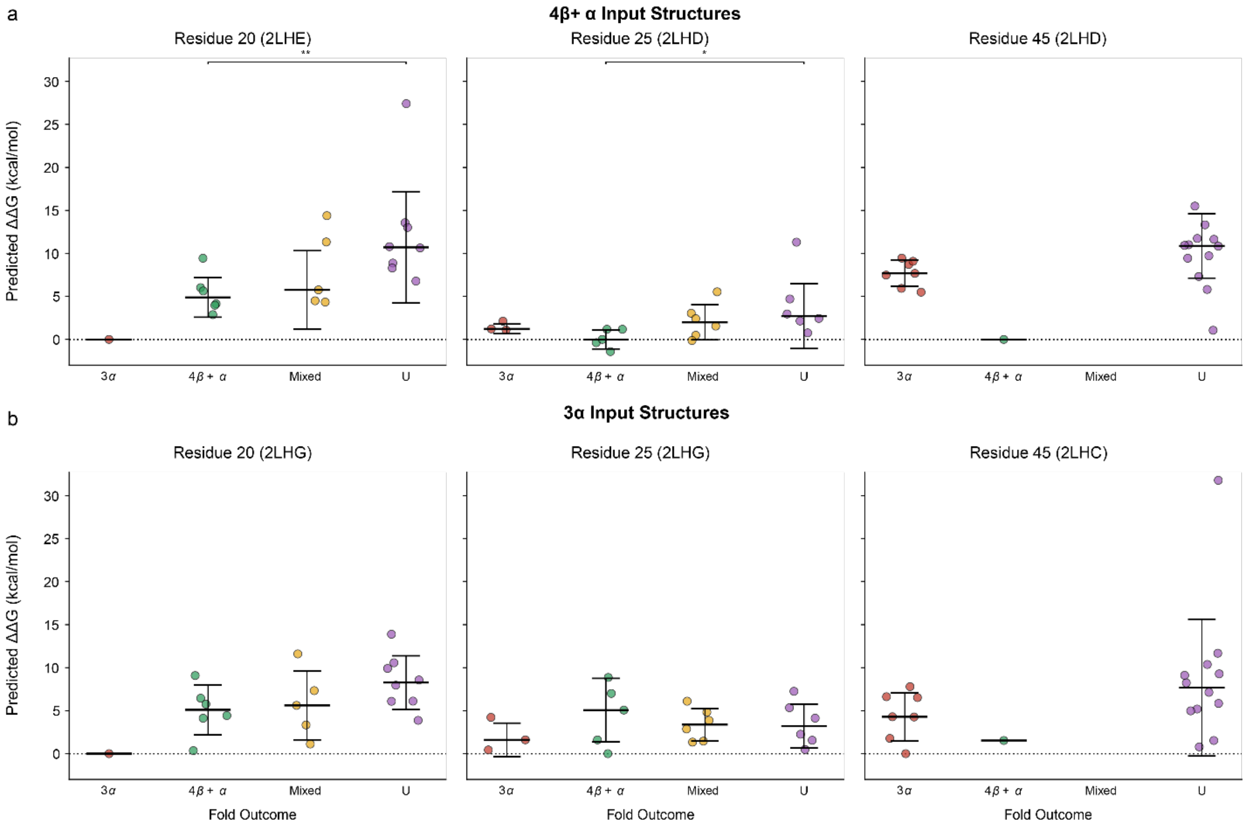


**Figure S5. Rosetta3 alternative protocol ΔΔG predictions of GA/GB point mutants.** The Rosetta3 CartDDG-Ref2015 (47) method was used to produce ΔΔG predictions for all residue 20, 25, and 45 mutants with both **(a)** 4β+α and **(b)** 3α fold reference structures as input. Variants were classified as 3α, 4β+α, or unfolded (U) if that state accounted for at least 80% of the estimated fold state based on NMR measurements. If no single fold outcome met this threshold, then the variant was classified as "mixed". Two-sided Mann-Whitney U testing was used to perform an inter-group comparison between all groups with more than one mutant, and statistically significant comparisons (* $p < 0.05$, ** $p < 0.01$, *** $p < 0.001$) are indicated above the groups.

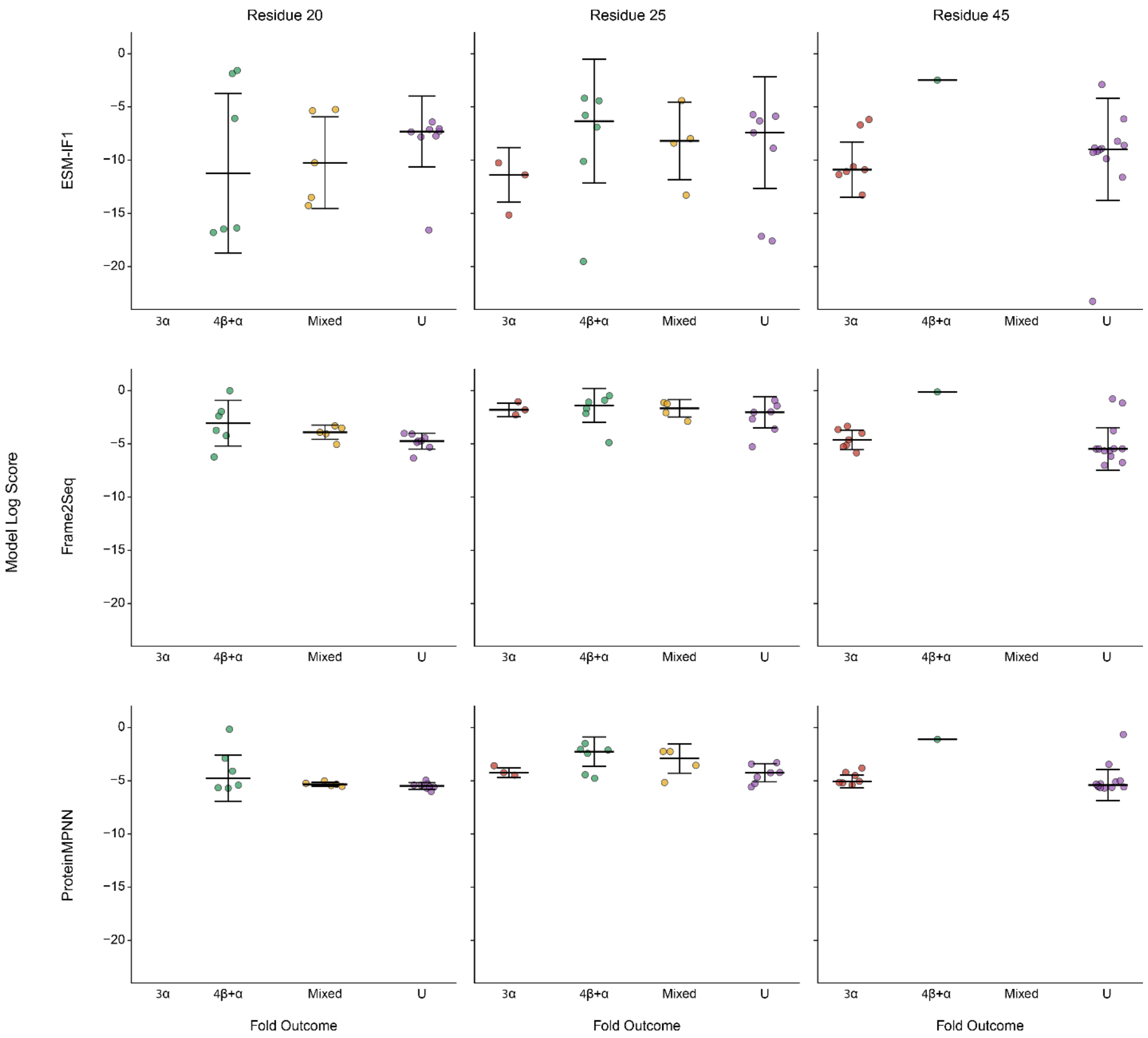


**Figure S6. Inverse folding method scores for 4β+α inputs structures.** Variants were grouped according to fold outcome as determined by NMR. ESM-IF1 (44), Frame2Seq (45), and ProteinMPNN (43) were used to score the respective 4β+α structures for variants at each residue position. Two-sided Mann-Whitney U test was used to determine significance of each group pairing, with only significant relationships annotated (* $p < 0.05$, ** $p < 0.01$, *** $p < 0.001$).

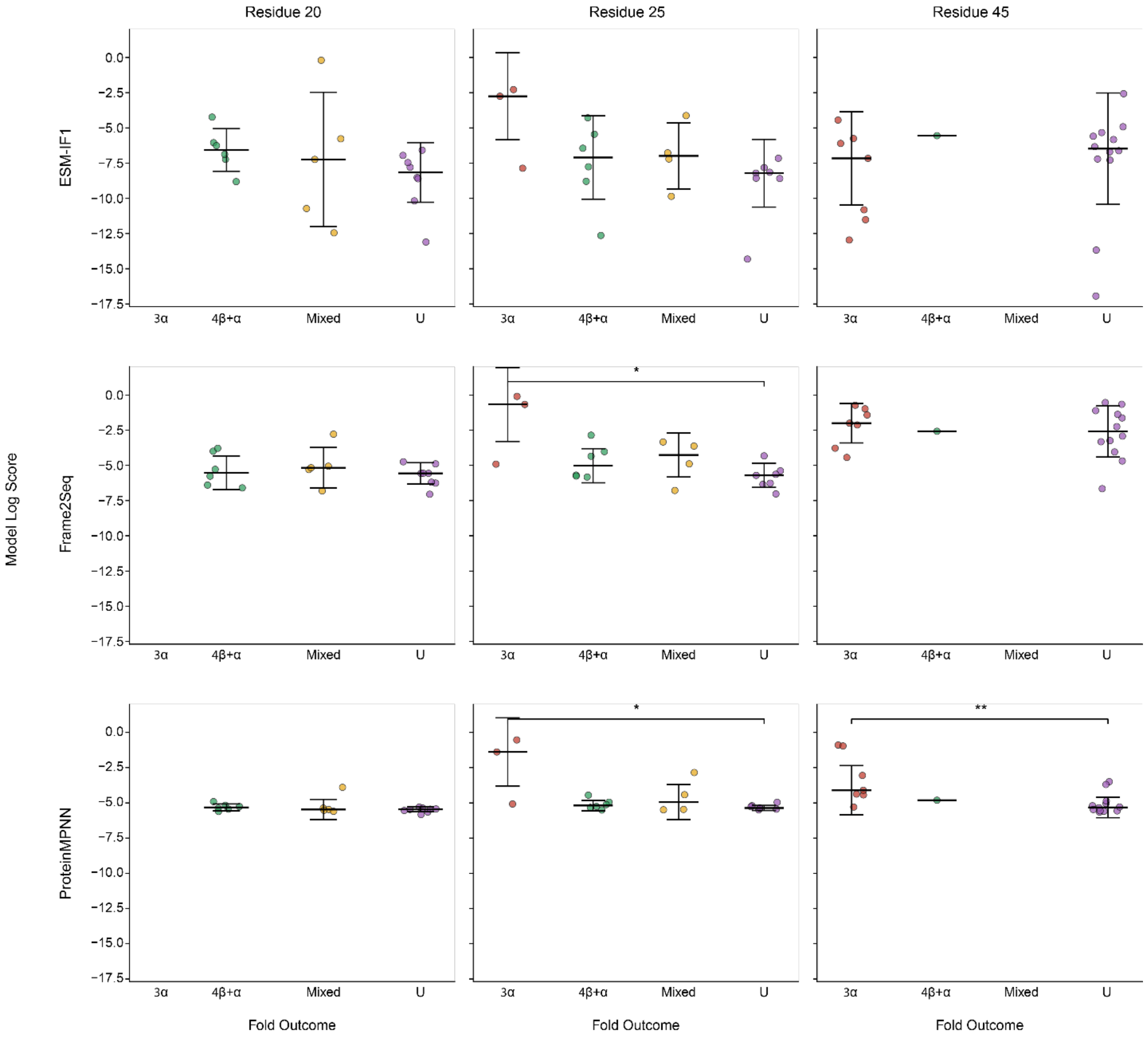


**Figure S7. Inverse folding method scores for 3α input structures.** Variants were grouped according to fold outcome as determined by NMR. ESM-IF1, Frame2Seq, and ProteinMPNN were used to score the respective 3α structures for variants at each residue position. Two-sided Mann-Whitney U test was used to determine significance of each group pairing, with only significant relationships annotated. (* $p < 0.05$, ** $p < 0.01$, *** $p < 0.001$)

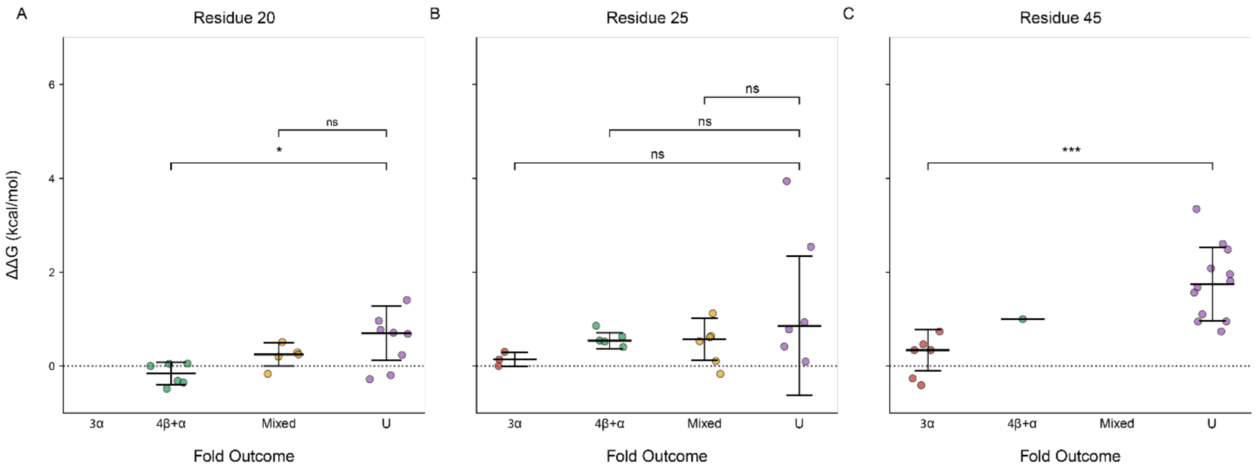


**Figure S8. Comparison of individual residue variant groups and MegaScale thermostability measurements.** Variants were grouped according to fold outcome as determined by NMR and experimentally determined ΔΔG values for each variant were parsed out of the MegaScale dataset (49). ΔΔG values of each folded group were compared to the unfolded group using a two-side Mann-Whitney U test, with significance or lack of significance (ns) annotated above the pairs.

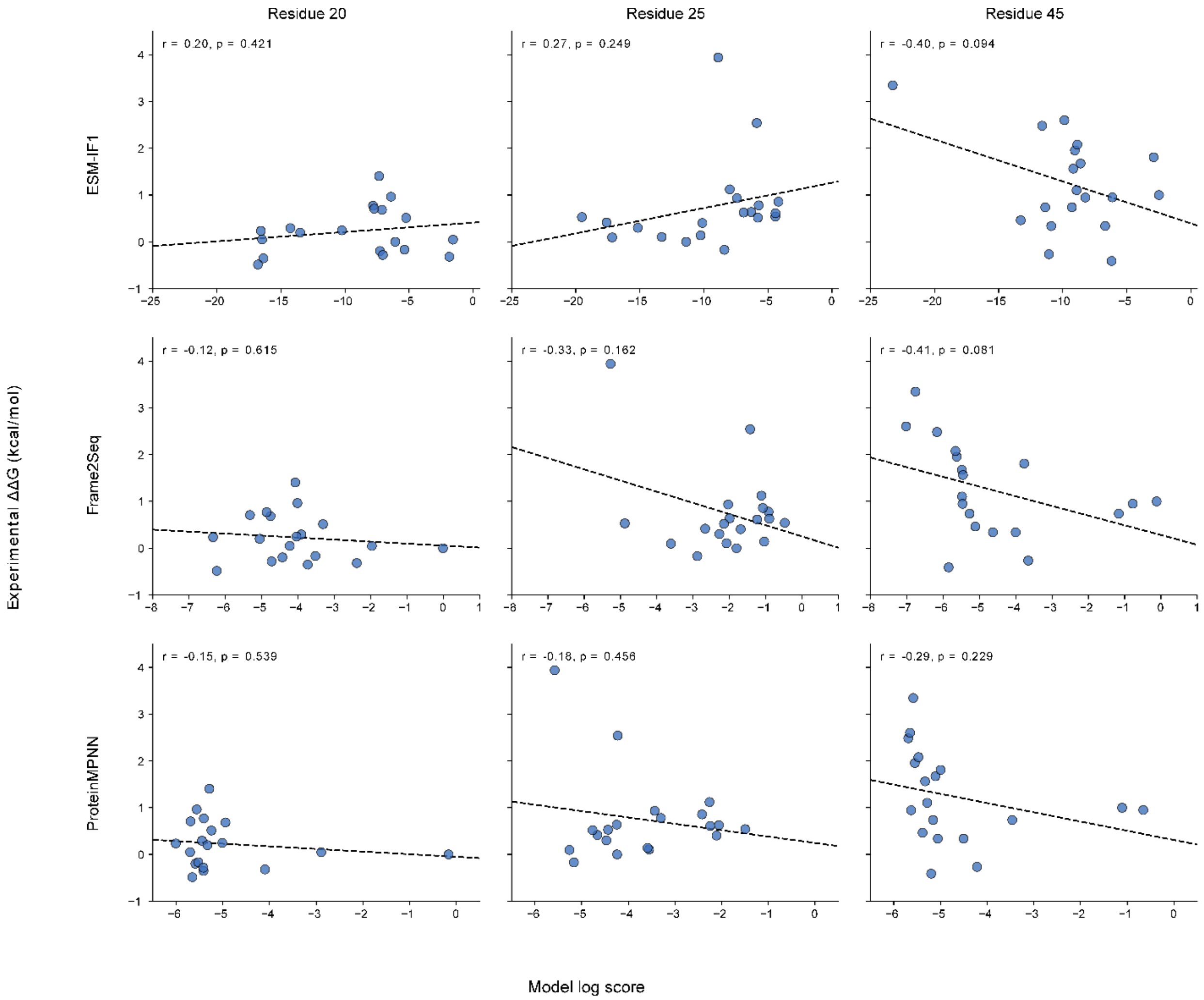


**Figure S9. Correlation of inverse folding scores of 4β+α input structures and ΔΔG measurements.** ESM-IF1, Frame2Seq, and ProteinMPNN were used to score the respective 4β+α structure for variants at each residue position. These values were plotted against their respective experimentally determined ΔΔG values parsed from the MegaScale dataset. Pearson correlations were calculated, with significance determined via two-sided correlation t-test.

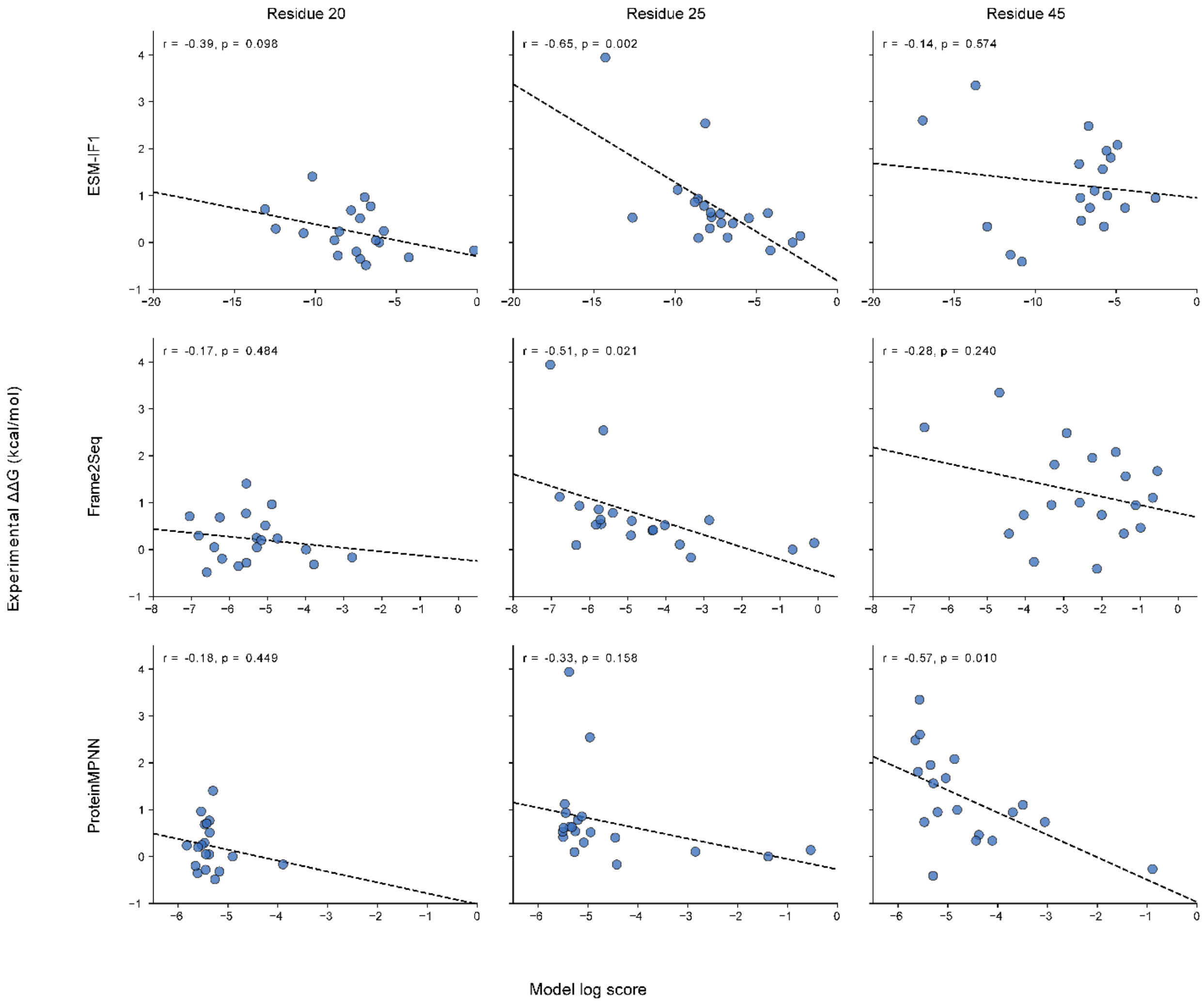


**Figure S10. Correlation of inverse folding scores of 3α input structures and ΔΔG measurements.** ESM-IF1, Frame2Seq, and ProteinMPNN were used to score the respective 3α structure for variants at each residue position. These values were plotted against their respective experimentally determined ΔΔG values parsed from the MegaScale dataset. Pearson correlations were calculated, with significance determined via two-sided correlation t-test.

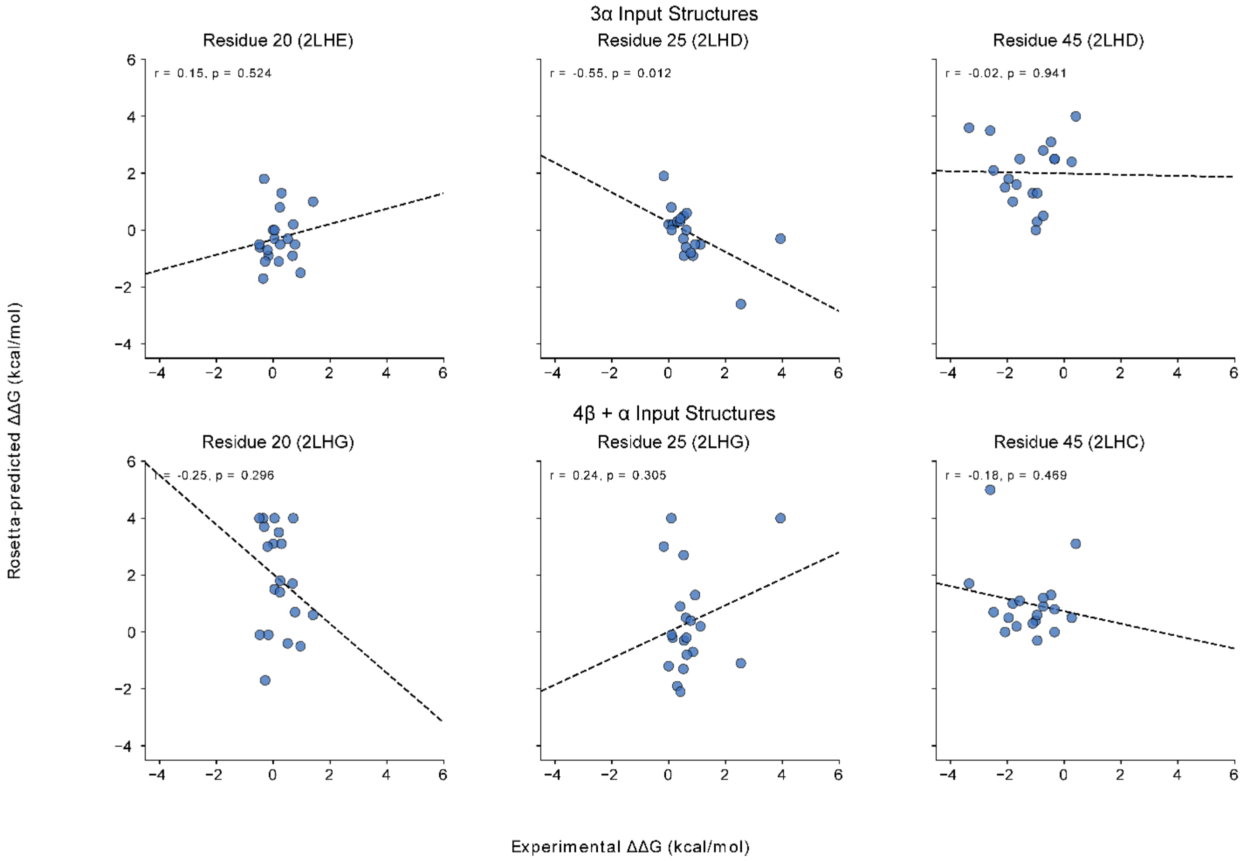


**Figure S11. Correlation of Rosetta ΔΔG and MegaScale experimental ΔΔG measurements.** Rosetta2.3 monomer ΔΔG prediction protocol (46) was used to predict ΔΔG values for variants at each residue position using both their respective 4β+α and 3α structures as input. Both sets of these values were plotted against their respective experimentally determined ΔΔG values parsed from the MegaScale dataset. Pearson correlations were calculated, with significance determined via two-sided correlation t-test.

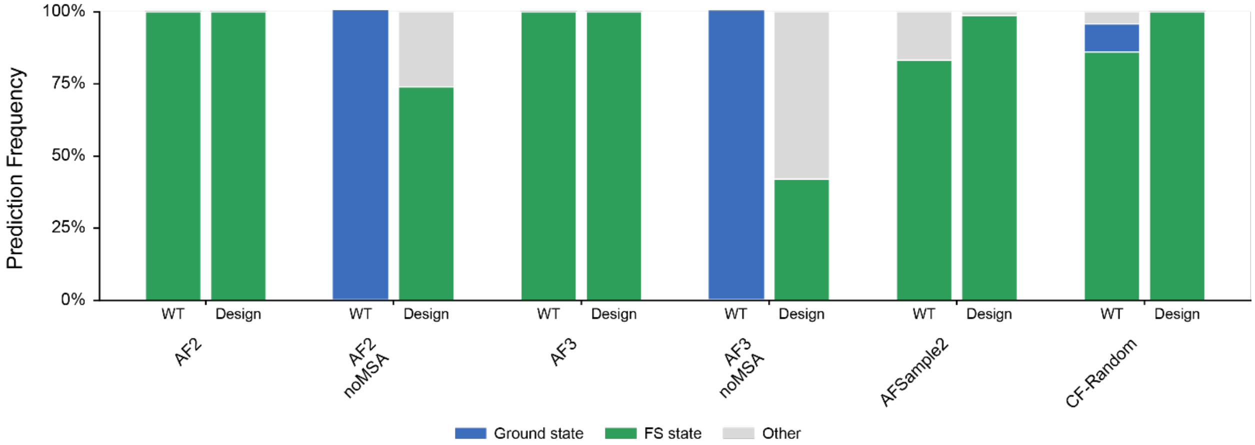


**Figure S12. Modeling of KaiB wild type and designed sequences.** AlphaFold2 (18), AFSample2 (24), AlphaFold3 (19), and CF-Random (25) were used to produce 1000 models for the wild type sequence of the KaiB metamorphic protein, as well as the designed KaiB described by Wayment-Steele et al (23). Model output was classified as being in either the ground state, fold switch state, or neither based on alignment to reference structures 2QKE and 5JYT, respectively. As the design is based on AF-Cluster modeling, AF-Cluster was omitted from the evaluation. Wild type outcomes (left bars) are expected to be 89% ground state, while the designed sequence is expected to be majority fold switch state (FS, 86%), based on previous NMR characterization (23).

| Sequence | % Identity to GB98 | Avg TM Score 3α | Avg TM Score 4β+α | Avg Confidence | Num Models 3α | Num Models 4β+α | Num Models Unfolded | % Models 3α | % Models 4β+α | % Models Unfolded |
|---|---|---|---|---|---|---|---|---|---|---|
| TTYKLILNLKQAKEEA**I**KELVDAG**I**AEK Y**I**KLIANAKTVEGVWTYKDEI**LKA**TVTE | 88 | 0.63 | 0.42 | 81.68 | 1000 | 0 | 0 | 100 | 0 | 0 |
| TTYKLILNLKQAKEEATKELVDAGTAEK Y**I**KLIANAKTVEGVWTYKDEI**LKA**TVTE | 92 | 0.63 | 0.41 | 81.87 | 1000 | 0 | 0 | 100 | 0 | 0 |

**Table S1. AlphaFold2 modeling results for GA/GB homologs with Tyr at position 45.** Two GA/GB mutant sequences previously assessed to fold into the 3α state with a tyrosine present at position 45 were predicted using AlphaFold2. Models were compared to reference GA structure 2LHC and reference GB structure 2LHD using TM-align, with models being classified as 3α or 4β+α if the TM-score of the alignment against one of the structures was greater than 0.5. Position 45 is underlined and colored red. Positions which deviate from their identity in GB sequence are bolded and underlined.

| PDB_ID | TMScore_2LHC_3A | TMScore_2LHD_4b+a | % Identity | E_Value |
|---|---|---|---|---|
| 1EM7 | 0.398 | 0.699 | 58.9 | 6.08E-11 |
| 1FCC | 0.362 | 0.362 | 58.9 | 4.74E-12 |
| 1IGC | 0.303 | 0.401 | 58.9 | 6.53E-12 |
| 1MHX | 0.402 | 0.717 | 50.9 | 9.38E-08 |
| 1MI0 | 0.397 | 0.725 | 52.7 | 1.90E-08 |
| 1MVK | 0.296 | 0.337 | 53.7 | 3.60E-08 |
| 1PGB | 0.395 | 0.700 | 61.8 | 8.97E-12 |
| 1PGX | 0.403 | 0.694 | 58.9 | 6.53E-12 |
| 1QKZ | 0.389 | 0.682 | 57.1 | 1.70E-11 |
| 1TF0 | 0.461 | 0.458 | 59.5 | 2.04E-04 |
| 1UWX | 0.392 | 0.683 | 57.1 | 1.70E-11 |
| 2J5Y | 0.622 | 0.423 | 59.5 | 0.0002036 |
| 2ON8 | 0.403 | 0.692 | 53.7 | 7.29E-09 |
| 2ONQ | 0.394 | 0.706 | 53.7 | 2.03E-09 |
| 2QMT | 0.402 | 0.696 | 61.1 | 4.42E-11 |
| 2VDB | 0.412 | 0.423 | 59.5 | 0.0002036 |
| 2ZW0 | 0.402 | 0.708 | 61.8 | 1.23E-11 |
| 2ZW1 | 0.388 | 0.720 | 61.8 | 4.42E-11 |
| 3FIL | 0.385 | 0.667 | 59.2 | 2.03E-09 |
| 3MP9 | 0.402 | 0.682 | 61.8 | 8.974E-12 |
| 3U4E | 0.400 | 0.737 | 57.8 | 8.26E-06 |
| 3UI3 | 0.374 | 0.586 | 61.1 | 4.42E-11 |
| 3V3X | 0.389 | 0.544 | 57.4 | 1.00E-08 |
| 4DQO | 0.341 | 0.355 | 57.8 | 8.26E-06 |
| 4KGR | 0.293 | 0.649 | 61.8 | 8.97E-12 |
| 4KGS | 0.383 | 0.660 | 60 | 4.42E-11 |
| 4KGT | 0.383 | 0.694 | 56.3 | 1.29E-07 |
| 4OZA | 0.315 | 0.680 | 61.8 | 6.53E-12 |
| 4OZB | 0.311 | 0.661 | 60 | 1.15E-10 |
| 4OZC | 0.316 | 0.640 | 58.1 | 1.48E-09 |
| 4Q0C | 0.357 | 0.363 | 59.2 | 2.18E-10 |
| 4WH4 | 0.388 | 0.694 | 55.5 | 1.90E-08 |
| 5BMG | 0.394 | 0.681 | 59.2 | 7.80E-10 |
| 5BMI | 0.398 | 0.690 | 59.2 | 3.00E-10 |
| 5HFY | 0.307 | 0.670 | 61.8 | 1.23E-11 |
| 5HG2 | 0.309 | 0.656 | 60 | 8.36E-11 |
| 5HI1 | 0.287 | 0.645 | 60 | 1.15E-10 |
| 5LDE | 0.472 | 0.551 | 61.1 | 4.42E-11 |
| 5O94 | 0.393 | 0.753 | 50 | 1.21E-06 |
| 6C9O | 0.389 | 0.688 | 61.1 | 4.42E-11 |
| 6CHE | 0.377 | 0.671 | 59.2 | 2.18E-10 |
| 6CMX | 0.407 | 0.407 | 50.9 | 3.37E-07 |

| | | | | |
|---|---|---|---|---|
| 6CNE | 0.400 | 0.685 | 59.2 | 8.36E-11 |
| 6CPZ | 0.401 | 0.691 | 59.2 | 1.15E-10 |
| 6CTE | 0.387 | 0.691 | 59.2 | 1.15E-10 |
| 6KMC | 0.388 | 0.712 | 60 | 4.12E-10 |
| 6L91 | 0.403 | 0.719 | 60 | 8.36E-11 |
| 6L9B | 0.396 | 0.718 | 60 | 8.36E-11 |
| 6L9D | 0.396 | 0.696 | 60 | 6.08E-11 |
| 6LJI | 0.395 | 0.715 | 60 | 1.15E-10 |
| 6NL6 | 0.380 | 0.715 | 52.7 | 4.95E-08 |
| 6NL7 | 0.402 | 0.715 | 54.5 | 7.29E-09 |
| 6NL9 | 0.395 | 0.673 | 61.8 | 2.34E-11 |
| 6NLB | 0.396 | 0.707 | 60 | 4.12E-10 |
| 6O41 | 0.331 | 0.432 | 58.9 | 6.53E-12 |
| 6OC7 | 0.306 | 0.425 | 58.9 | 6.53E-12 |
| 6U8C | 0.398 | 0.709 | 50 | 2.61E-08 |
| 6UUH | 0.329 | 0.403 | 58.9 | 6.53E-12 |
| 6UYG | 0.293 | 0.398 | 58.9 | 6.53E-12 |
| 6V9I | 0.444 | 0.435 | 61.1 | 4.42E-11 |
| 6WAS | 0.347 | 0.416 | 57.8 | 8.26E-06 |
| 6WIT | 0.357 | 0.434 | 57.8 | 8.26E-06 |
| 6XSN | 0.357 | 0.333 | 56.7 | 4.10E-05 |

**Table S2. PDB structures of GA/GB-related proteins in possible training sets of AlphaFold and related deep learning methods**. The GA98 protein sequence was searched against the PDB database to compile a list of structures with at least 50% sequence identity resolved through cryo-EM or X-ray crystallography released before 09-30-21. Each structure was assessed for structural similarity (TM-score) against 3α and 4β+α structures, and sequence identity and BLAST E-value from are also shown for reference.
